# Loss of TET2 activity limits the ability of vitamin C to activate DNA demethylation in human HAP1 cells

**DOI:** 10.1101/2025.06.27.661936

**Authors:** Maciej Gawronski, Marta Starczak, Aleksandra Wasilow, Tomasz Dziaman, Ryszard Olinski, Daniel Gackowski

## Abstract

**Background:** The TET family of proteins - TET1, TET2, and TET3 - are α-KG and Fe^2+^ dependent dioxygenases that play crucial roles in active DNA demethylation and the deposition of epigenetic marks such as 5-hydroxymethylcytosine, 5-formylcytosine, and 5-carboxycytosine. TET proteins can also oxidize thymine to 5-hydroxymethyl uracil - a modification whose role is still poorly understood. TET proteins add a new layer of information in regulating gene expression, cellular development, and lineage specification. Dysregulation of TET activity is implicated in various cancers, especially in hematological malignancies, where TET2 loss-of-function mutations are prevalent. TET2’s role in hematopoiesis is critical, as its knockdown skews progenitor differentiation toward the myeloid lineage and drives carcinogenesis. Therefore, restoring the lost activity of TET proteins is often proposed as an important component of cancer treatment.

This study explores the distinct contributions of TET paralogs in generating active demethylation products in malignant cells. It examines whether vitamin C, a known cofactor of many dioxygenases, can compensate for the loss of specific TET paralogs. We applied a highly sensitive and specific methodology (2D-UPLC-MS/MS) to assess TET activity in the HAP1 cell line with single and double TET functional knockouts and in cells with the activity of all TET proteins impaired.

**Results:** Our findings reveal that TET2 is essential for all steps of iterative oxidation, and its loss has the most significant effect on 5-hydroxymethylcytosine and 5-formylcytosine levels. Vitamin C enhances TET activity and increases the levels of these oxidation products. However, its effect in TET2 knockout cells is limited; Vitamin C increased cytosine modification levels in TET2KO cells, but not to the extent observed in treated wild-type cells, indicating incomplete compensation for TET2 loss.

**Conclusions:** Our results demonstrated that each TET protein has a distinct, separate contribution to generating active demethylation products. The absence of individual TET paralog is linked with the specific pattern of active demethylation products in DNA, which is preserved after vitamin C treatment. Therefore, the deletion of one of the TET enzymes cannot be compensated for by the increased activity of the other TET family members, highlighting the unique roles of each TET paralog in epigenetic regulation.

## BACKGRROUND

In 2009, two seminal studies reported the enzymatic oxidation of 5-methylcytosine (5-mCyt) to 5-hydroxymethylcytosine (5-hmCyt) by proteins from the TET (ten-eleven translocation) family (1, 2). To date, three TET proteins have been identified in humans: TET1, TET2, and TET3. These enzymes catalyze the oxidation of 5-mCyt to 5-hmCyt, followed by further oxidation to 5-formylcytosine (5-fCyt) and ultimately 5-carboxycytosine (5-caCyt) (3). The latter two products are recognized and excised by glycosylases of the base excision repair pathway, such as thymine DNA glycosylase, leading to their replacement with unmodified cytosine (4, 5). Moreover, some studies suggest a sequential mechanism in which TET-mediated oxidation proceeds through each intermediate without the enzyme dissociating from DNA, until the final product, 5-caCyt, is formed (6). Others propose that the enzyme may dissociate from DNA after each oxidative step (7).

Additionally, TET proteins exhibit enzymatic activity towards thymine, oxidizing it to 5-hydroxymethyluracil (5-hmUra) (8). Depending on the pathway of formation, 5-hmUra can either form mutagenic 5-hmUra mispairs, following deamination of 5-hmCyt, or act as a potential epigenetic marker when formed via thymine oxidation, leading to stable 5-hmUra-adenine pairs (9).

The three human TET proteins exhibit differences in genomic localization, structure, and tissue-specific expression levels, implying both overlapping and distinct functional roles (10). Structurally, TET1 and TET3 contain an N-terminal CXXC zinc finger domain responsible for DNA binding, particularly within CpG dinucleotides, recognizing methylated and hydroxymethylated DNA (11). However, in neurons and neuronal progenitor cells, alternative promoter usage and splicing favor expression of a short TET3 isoform (Tet3s) that lacks the N-terminal CXXC domain. Consequently, Tet3s constitutes the majority of TET3 protein in these cell types. This structural difference has functional implications for DNA-binding properties and gene targeting (12). Similarly to Tet3s, TET2 has also lost CXXC domain, which now exists as a separate protein, IDAX/CXXC4, regulating TET2 activity and facilitating its caspase-dependent degradation (13). Moreover, TET1 contains nuclear localization sequences and regions responsible for interacting with thymine DNA glycosylase (2, 14, 15).

The expression of TET proteins is tightly regulated through alternative splicing, promoter usage, and post-transcriptional modifications, generating several isoforms with tissue-specific functions, some of which extend beyond their catalytic activity (16–19). These isoforms may contribute to tissue-specific demethylation processes and gene regulation, preventing de novo methylation by anchoring to unmethylated regions of DNA (20, 21).

TET3 is highly expressed in oocytes and zygotes, where it plays a role in the global demethylation of the paternal genome and partial demethylation of maternal DNA, resetting the epigenetic landscape to facilitate totipotency in the zygote and early embryonic cells (22–25). TET1 helps regulate pluripotency genes such as NANOG and also represses differentiation-related genes, potentially preventing de novo methylation at key CpG islands in pluripotency gene promoters (26, 27).

TET2 plays a critical role in hematopoiesis. Its deficiency results in impaired differentiation of hematopoietic stem cells, promoting the expansion of the monocyte/macrophage lineage (28). Knockout models of TET2 in mice show uncontrolled hematopoietic cell proliferation and the development of malignancies resembling human hematologic disorders, such as chronic myelomonocytic leukemia and myeloid leukemias (29). In human CD34+ cord blood cells, TET2 deficiency similarly skews differentiation toward the granulocyte/monocyte lineage, at the expense of lymphoid and erythroid cells (30). Mutations in the TET2 gene have been identified in myelodysplastic syndromes and myeloproliferative disorders (31, 32), as well as in chronic myelomonocytic leukemia and acute myeloid leukemia (10, 33, 34), and are linked to poor prognosis and increased risk of disease progression (10). Mutations in TET2 are also found in a significant proportion of lymphomas, including 12% of diffuse large B-cell lymphoma cases and nearly half of angioimmunoblastic T-cell lymphomas (35, 36).

Vitamin C, a known cofactor for iron- and α-ketoglutarate-dependent dioxygenases, stimulates TET protein activity, by maintaining the catalytic iron in their active site in its Fe(II) state (37). Studies have shown that adding vitamin C to embryonic stem cell cultures increases 5-hmCyt levels, especially in promoter regions, without affecting TET gene expression, suggesting a direct effect on enzyme activity (38–40). Vitamin C is believed to enhance the catalytic activity of TET proteins by reducing Fe(III) to Fe(II) in the enzyme’s active site, restoring its function and accelerating 5-mCyt oxidation (41).

Furthermore, vitamin C’s antioxidant properties may protect the catalytic iron properties, preserving enzyme activity (42). Results of studies conducted on bladder cancer cells by Peng et al. showed that vitamin C increased genomic 5-hmCyt in a time- and concentration-dependent manner and did not change TET transcript levels, indicating it boosted enzyme activity. At higher doses (0.5–5 mM), vitamin C inhibited cancer cell growth and induced apoptosis(43). Agathocleous et al. found that mouse and human hematopoietic stem cells (HSCs) naturally accumulate high ascorbate. Moreover, systemic vitamin C depletion (genetically or by diet) increased HSC self-renewal by reducing Tet2 activity, and cooperated with oncogenic FLT3-ITD to accelerate leukemia. Those effects were reversed by dietary supplementation of vitamin C (44). Recent findings suggest that in cases of TET2 deficiency, vitamin C supplementation can restore TET activity, increase global levels of oxidized 5-mCyt products, and delay the progression of hematological malignancies (45). Brabson et al. demonstrated that vitamin C enhances TET2 activity in acute myeloid leukemia (AML), promoting DNA demethylation and myeloid differentiation. When combined with PARP inhibitors, vitamin C induces synthetic lethality in AML cells by increasing oxidized methylcytosine levels and trapping PARP1 at sites of replicative stress (46). In light of the above findings, and particularly considering the potential use of vitamin C as an adjuvant in current therapeutic strategies, a precise understanding of its impact on the activity of individual TET enzymes is of significant biological and clinical relevance.

## METHODS

### Cell lines and cell culture

All experiments involving the use of cell cultures were performed in sterile conditions, maintaining aseptic principles following Good Laboratory Practices in a laboratory meeting the requirements of biological safety class II.

HAP1 line - an adherent, haploid cell line derived from the KBM-7 chronic myeloid leukemia cell line, established from the cells of a 40-year-old man, which, through genetic modifications, lost the second copy of chromosome 8 of the parent cell line and also retained a 30-Mbp fragment of chromosome 15, connected to chromosome 19.

For the conducted research, the parent HAP1 line (Horizon Discovery, cat. no. c631) was purchased from Horizon Discovery, as well as commercially available variants of this line in which functional gene knockouts were introduced using the CRISPR/Cas9 technique:

- TET1 (20 bp deletion in exon 2, Horizon Discovery, cat. no.: HZGHC001121c002);
- TET2 (8-bp deletion in exon 3, Horizon Discovery, cat. no.: HZGHC003562c011);
- TET3 (14-bp deletion in exon 4, Horizon Discovery, cat. no. HZGHC001112c004);

These lines constitute a model of hematologic cancer with a deactivating mutation in one of the proteins of the TET protein family.

Horizon Discovery was also commissioned to create three unique line variants using the CRISPR/Cas9 technique, with functional pair knockouts:

- TET1/TET2 (deletion of 20 bp in exon 2 of the TET1 gene/deletion of 26 bp in exon 3 of the TET2 gene; Horizon Discovery, cat. no.: HZGHC007806c011);
- TET1/TET3 (1 bp insertion in exon 2 of the TET1 gene/1 bp deletion in exon 4 of the TET3 gene; Horizon Discovery, cat. no. HZGHC007804c010);
- TET2/TET3 (deletion of 14 bp in exon 3 of the TET2 gene/deletion of 14 bp in exon 4 of the TET3 gene; Horizon discovery, cat. no. HZGHC007813c001);

The cells of these lines constitute a model in which only one of the TET family proteins remains active, which enables the study of the enzymatic activity of individual TET proteins and the potential impact of the tested modulators on a specific enzyme. It is also worth noting that mRNA and protein can still be produced in cell lines with a functional knockout of the TET genes.

Cell cultures of all HAP1 line variants were performed based on Horizon Discovery guidelines.

A complete culture medium was prepared for each of the cultivated lines, consisting of 10% Fetal Bovine Serum (FBS Good Forte Filtrated Bovine serum EU approved, PAN Biotech, cat. no.: P40-47500), 1% antibiotic solution (Antibiotic Antimycotic Solution, 100X, Corning, cat. no.: 30-004-CI) and Iscove’s Modification of DMEM (Corning, cat. no.: 10-016-CV), selected based on the recommendations of the manufacturer, ensuring optimal cell growth.

Cells of each of the tested lines were seeded into culture bottles with an area of 75 cm^2^ (Nunc™ EasYFlask™ 75cm^2^ Cell Culture Flasks, Thermo-Fisher cat. no.: 156499), containing 10 ml of complete culture medium.

After 24 hours, the culture medium was poured off, and the vessels were rinsed twice with a warm Dulbeccòs Phosphate Buffered Saline solution, thus removing dead cells and those not stuck to the bottom of the culture vessel; then, 10 ml of fresh medium was added. Cultures were carried out in a sterile incubator at 37 °C and in an atmosphere containing 5 % CO_2_. After the culture time set in each experiment had elapsed, the culture medium was removed from the culture vessels, the cells were washed twice with warm Dulbeccòs Phosphate Buffered Saline solution, and then they were collected by trypsinization. The obtained cell pellet was transferred to a low-temperature freezer and stored at - 80 °C until the genetic material isolation procedure was performed.

### Vitamin C experiments

All cell lines were grown in accordance to the methodology above. During all vitamin C experiments ascorbic acid from Sigma (cat. no. A44544-100G) was used. Briefly, HAP1 cells (with single or double knock-outs) were grown to 50 % confluence. Then, cell culture medium was discarded, cells were rinsed with warm PBS, and fresh medium containing 100 µmol/l of vitamin C was added (100 milimol/l vitamin C stock solution was freshly prepared, and timidity was added to the medium to achieve a 100 µmol/l concentration.). The concentration of L-ascorbic acid (vitamin C) was selected to reflect the high plasma levels of this compound observed in healthy, well-nourished individuals, reported in the literature, as well as based on previously conducted MTT tests, in which the effect of 24-hour exposure of HAP1 cells to 1 µmol/l, 10 µmol/l, 100 µmol/l and 1 mmol/l of vitamin C was analyzed (No significant differences were observed for the first three concentrations, while the concentration of 1 mmol/l proved cytotoxic.

The results of the MTT test are presented in Supplementary Figure 1 in Additional file 2). Control cells were cultured without vitamin C. After 24 hours of incubation cells were harvested and DNA was isolated from them, by method described below.

### siRNA experiments

HAP1 TET1/TET3KO cells were used in siRNA experiments. By inhibition of TET2 expression in this line we obtained a model in which almost no TET protein activity is present, simulating triple knockout. 3 unique 27mer anti-TET2 siRNA duplexes (cat. No.: SR30002), Universal scrambled negative control siRNA duplex (cat. no.: SR30004) as well as siTran 2.0 transfection reagent (cat. no. TT320001) were bought from OriGene. All siRNA experiments were conducted in accordance with manufacturers protocols. siRNA concentration, transfection time, and volume of transfection reagent were optimized before performing final experiments. By using the manufacturer’s approach of simultaneously using three different siRNA duplexes, we achieved (in HAP1 TET1KO/TET3KO cells) an average reduction of TET2 mRNA expression of 74% during 24-48h transfection time (see Supplementary Figure 2 in Additional File 2), which is consistent with the manufacturer’s guarantee of >70% expression reduction. Briefly, the procedure for conducting the final experiment was as follows: cells were seeded (in Iscove’s Modification of DMEM with 10 % Fetal Bovine Serum) on T25 flask 24 hours before transfection in a density (0.8 x 10^6^ cells per flask) that enables achieving 50 % confluence at the time of transfection. Then, cell culture medium was discarded, cells were rinsed with warm PBS (3 ml), and fresh culture medium (5 ml) was added. Diluted combination of three siRNA duplexes or scrambled negative control siRNA were mixed with siTran 2.0 (4.2 µl of each duplex, 300 µl 1X transfection buffer and 7.2 µl siTran 2.0). Mixture was incubated for 15 minutes at room temperature and subsequently added dropwise to cell culture to achieve 25 nmol/l of each duplex. Moreover, to some part of transfected cells vitamin C in 100 µmol/l concentration was added, and control cells were grown without any additives. Transfection lasted for 24 hours, during which the cells were kept in a CO_2_ incubator at 37 ⁰C and in atmosphere containing 5 % CO_2_. After 24 hours all cells were harvested and DNA was isolated from them, by method described below.

### DNA isolation from cell culture samples

This work uses the method of determining epigenetic modifications in DNA using two-dimensional, ultra-performance liquid chromatography with tandem mass detection developed and optimized at Department of Clinical Biochemistry (Nicolaus Copernicus university in Toruń, Ludwik Rydygier Collegium Medicum in Bydgoszcz), based on a previously published method (47), which was subject to significant modifications and described in detail in the monograph entitled “DNA Modifications Methods and Protocols” (48). The assumptions of this method are briefly presented below.

The frozen cell pellets were brought to room temperature. After thawing, the remains of the culture medium were pipetted out and then suspended in buffer “B” (5 mmol/l Na_2_EDTA, 0.15 mmol/l deferoxamine, 10 mmol/l Tris-HCL, pH= 8.0) in a volume ranging from 0.5 to 3 ml. The buffer volume was selected experimentally and depended on the type of cells and their number. An appropriate volume of RNAse A solution from bovine pancreas (1 mg/ml; Sigma-Aldrich, cat. no.: R6513-1G) was added to the cell suspension so that the final concentration in the sample was 30 µg/ml, and a solution of RNAse T1 from *Aspergillus oryzae* (1 U/µl; Sigma-Aldrich, cat. no.: R1003-500KU), to a final concentration in the sample of 10 U/ml to limit possible contamination of the isolated DNA with RNA molecules.

Cells were then lysed by adding an appropriate volume of 10% (m/v) Sodium Dodecyl Sulfate solution so that the final concentration of this compound in the sample was 0.5 %. The cell lysate prepared this way was incubated for 30 minutes in a water bath heated to 37 °C.

In order to enzymatically digest proteins, present in the cell, in particular proteins associated with DNA, an appropriate volume of Proteinase K solution from *Tritirachium album* (20 mg/ml, Sigma-Aldrich, cat. no.: SRE0005) was added to the cell lysate so that the final concentration of this compound in the lysate was 4 mg/ml, and then incubated for 90 minutes in a water bath heated to 37 °C.

To remove cellular proteins as well as other water-insoluble substances, an equal volume of phenol-chloroform-isoamyl alcohol mixture (prepared in a volume ratio of 25:24:1) was added to the cooled samples, shaken vigorously for several minutes, and then centrifuged for 15 minutes, at a temperature of 4 °C, with an acceleration of 2800 x g. After centrifugation, the aqueous phase was collected and transferred to new centrifuge tubes.

Then, an equal volume of the chloroform-isoamyl alcohol mixture (prepared in a volume ratio of 24:1) was added to the previously collected aqueous phase, shaken vigorously for several minutes, then centrifuged for 15 minutes at 4 °C at 2800 x g and collected again—aqueous phase into new centrifuge tubes.

In order to precipitate highly polymerized DNA, three volumes of cold, 96 % ethanol were added to the obtained aqueous phase. The samples were mixed by gently moving the test tube until long, white DNA strands appeared, which were pulled out using a plastic loop. The DNA obtained this way was transferred to a 0.5 ml Eppendorf tube and then hydrated by adding 50 µl of clean, sterile water, free from RNases and DNases. The hydrated DNA was then enzymatically hydrolyzed to individual nucleosides.

### Enzymatic DNA digestion

In order to obtain single deoxynucleotides, the following was added to each sample of hydrated DNA, as well as to a “blank” sample containing 50 µl of water: 50 µl of NP1 buffer (200 mmol/L ammonium acetate, 0.2 mmol/l ZnCl_2_, pH = 4.6), 1 µl of Nuclease P1 solution (1 U/µl, New England BioLabs® Inc.; cat. no.: M0660S) and 1 µl of tetrahydrouridine solution - cytidine deaminase inhibitor (10 mg/ml, Sigma-Aldrich, cat. no.: 584223 −25MG), mixed thoroughly and then incubated for three hours in a water bath heated to 37 °C.

Then, to remove hydrophilic phosphate groups from deoxynucleotides and obtain deoxynucleosides that can be separated by reversed-phase chromatography, 13 µl of 10% NH_4_OH and 6 µl of alkaline phosphatase (Thermolabile Recombinant Shrimp Alkaline Phosphatase (1,000 U/ml) were added to the hydrolyzate, New England BioLabs® Inc.; Cat. No.: M0371S).

The pH of the hydrolyzate was checked using a universal indicator, which should be at least 8.0. If necessary, the samples were alkalized with a small additional ammonium hydroxide. The hydrolysates were incubated for 90 minutes in a water bath heated to 37 °C, then transferred to filter plates (AcroPrep Advance 96-Well Filter Plates 10 K) and centrifuged for 90 minutes at 4 °C at an acceleration of 1800 x g, thanks to which proteins and potential contaminants larger than the pore size of the membrane were removed from the mixture.

The obtained filtrates (100 - 120 µl) were transferred to 1.5 ml Eppendorf tubes and concentrated in a vacuum concentrator to 20 µl, removing acetate and ammonium ions, which evaporate under reduced pressure.

Then, 5 µl of a mixture of stable isotope-labeled internal standards was added to the concentrated hydrolysates, containing:

- [D_3_]-5-(hydroxymethyl)-2’-deoxycytidine (Toronto Research Chemicals, Inc.; cat. no.: H946632);
- [^13^C_10_,^15^N_2_]-5-methyl-2’-deoxycytidine (synthesis at the Department of Clinical Biochemistry);
- [^13^C_10_,^15^N_2_]-5-formyl-2’-deoxycytidine (synthesis at the Department of Clinical Biochemistry);
- [^13^C_10_,^15^N_2_]-5-carboxy-2’-deoxycytidine (synthesis at the Department of Clinical Biochemistry);
- [^13^C_10_,^15^N_2_]-5-(hydroxymethyl)-2’-deoxyuridine (synthesis at the Department of Clinical Biochemistry);

The concentration of each of the above internal standards in the standard mixture was 250 fmol/µl, and the process of preparing commercially unavailable internal standards is described in detail in chapter 10 of the monograph “DNA Modifications Methods and Protocols” (49).

### Chromatographic analysis of deoxynucleosides

Chromatographic determinations were made using a system consisting of an ACQUITY UPLC PLUS I-Class, Waters® chromatograph: two gradient pumps, an isocratic pump, two automatic, programmable, two-position, six-port valves, a high-pressure tee, a photodiode-array detector (PDA detector), an ultrasensitive tandem quadrupole mass spectrometer (Xevo TQ-XS) and a set of chromatographic columns, which included:

- pre-column: CORTECS T3 VanGuard (Waters®), 5 mm x 2.1 mm, 1.6 μm.
- first dimension column: CORTECS T3 (Waters®), 150 mm x 3 mm, 1.6 µm.
- trapping column: XSelect CSH C18 (Waters®), 20mm x 3 mm, 3.5 μm.
- second dimension column: ACQUITY UPLC CSH C18 (Waters®), 100 mm x 2.1 mm, 1.7 μm.

For the needs of the analyses, several eluents were prepared based on LC/MS class reagents and ultrapure water, which constituted the mobile phases in the chromatographic system used and included:

- mobile phase A1: CH_3_COOH 0.05 % (v/v),
- mobile phase A2: CH_3_COOH 0.01 % (v/v),
- mobile phase B1: acetonitrile;
- mobile phase B2: methanol;
- mobile phase C1: ultrapure water;

The chromatographic system was configured in such a way that the mixture of compounds first went to the pre-column, the purpose of which was to remove all mechanical impurities and trap substances that could irreversibly bind to the chromatographic bed of subsequent columns. Then the deoxynucleoside mixture was separated on a first dimension column (flow 0.5 ml/min, gradient: from start to 1 min 99.3 % of phase A1 and 0.7 of phase B1, curve 6; up to 7 min 50 % of phase A1, 50 % of phase B1, curve 8; up to 7.2 min 50 % of phase A1, 50 % of phase B1, curve 6; from 7.21 min to 9 min 99.3 % of phase A1 and 0.7 % of phase B1, curve 6), from which individual analytes reached the MS/MS detector in specific time windows (5-hydroxymethyl-2’-deoxycytidine, 5-formyl-2’-deoxycytidine) either, after dilution with phase C1 in a 1:1 ratio, onto a trapping column, (5-carboxy-2’-deoxycytidine, 5-hydroxymethyl-2’-deoxyuridine), or were directed to the PDA detector (2’-deoxyguanosine, 2’-deoxycytidine, 2’-deoxythymidine, 2’-deoxyadenosine, 5-methyl-2’-deoxycytidine), where they were detected in UV light in the range of 260-280 nm. This approach made it possible not only to assess the concentration of dT, dA, and 5-mCyt in the tested sample but also to monitor the quality of the enzymatic DNA hydrolysis process (based on the presence of nucleotides and oligonucleotides) as well as to determine the degree of contamination of the RNA sample (based on the presence of ribonucleosides).

The fractions retained on the trapping column were redirected to the second dimension column (flow 0.35 ml/min, gradient: from start to 1.8 min 99 % of the A2 phase and 1 % of the B2 phase, curve 6; up to 3 min 91 % of the A2 phase and 9 % of phase B2, curve 6; up to 4.2 min 50 % of phase A2 and 50 %, curve 7; up to 6 min 50 % of phase A2 and 50 % of phase B2, curve 6; from 6.01 to 9 min 99 % phase A2 and 1 % of phase B2, curve 6), where they separated. The separated, modified nucleosides went to the ion source, where they underwent positive or negative ionization in the electrospray (ES) mode, creating daughter ions that went to the detector.

It is worth noting here that internal standards labeled with stable isotopes from the chromatographic side behave the same as their unlabeled counterparts and leave the chromatographic column at the same time, while their ionized forms have a different mass-to-charge ratio, which is the basis for quantitative analysis of the tested modifications.

Before performing chromatographic determinations, the chromatographic system was configured in terms of signal strength and calibrated:

For analyses under UV light - through external calibration, consisting in the creation of calibration curves based on the analysis of calibration solutions (5 injections, 2 µl each) for each of the compounds analyzed by this method, in the ranges of 0.00781 to 1 mmol/l (for dA, dT, dC, and dG) and in the range of 0.39 - 50 µmol/l for 5-mCyt; For MS/MS analyses - through internal calibration, based on the ratio of analyte surface areas to labeled internal standards.

All samples were always analyzed in three to five technical replicates (injections), and the resulting chromatograms were quantitatively analyzed using TargetLynx™ software (Waters). Example chromatograms of “blanks” and samples are provided in Additional File 2 in Supplementary Figure 3.

Subsequently, the mean, standard deviation (SD), and relative standard deviation (RSD) were calculated from all technical replicates for a given sample, both native and modified deoxynucleosides. Results with RSD > 2.5 % for native deoxynucleosides and with RSD > 15 % for modified deoxynucleosides were rejected. The total concentration of nucleosides in the sample (dN), originating from double-stranded DNA, was calculated based on twice the sum of the concentrations of dT and dA, which can be written as dN = 2 x (dT + dA).

Relative amount of modified nucleosides content in the tested material was calculated by dividing their concentrations by the total concentration of deoxynucleosides in the sample.

### Statistical analysis of results

The results of all experiments were prepared in both graphical and tabular form in a Microsoft Excel 2019 spreadsheet and in Statistica 13.3 (Tibco Software Inc. (2017)).

The Excel file was used as a batch file for Statistica 13.3 (TIBCO Software Inc. (2017). Statistica, in which:

- the type of individual variables was determined; they were given codes and text labels,
- histograms were drawn for numerical variables for each cell line and each experiment,
- outlier results were removed based on the analysis of the generated histograms,

The significance of the results of individual experiments was analyzed using a one-way, two-way ANOVA test with post-hoc tests: Tukey’s HSD and Fisher’s NIR - the type of test was selected based on the analysis of variance. The cut-off value indicating statistical significance was p<0.05.

It was also decided to normalize the data obtained from experiments performed on HAP1 cells with single and double TET functional knockouts and show them as a fold change of observed genomic levels of epigenetic DNA modifications relative to their levels observed in control cells. For this purpose, the values observed in all cells cultured without vitamin C supplementation were divided by the mean value observed in untreated Parental HAP1 cells, as well as the values observed in all cells cultured with vitamin C were divided by the mean value observed in Parental HAP1 cells exposed to vitamin C.

## RESULTS

### Effect of vitamin C supplementation in HAP1 cell line with single knock-outs of TET genes

To determine the effect of mutations in a single gene encoding individual TET proteins on the genomic content of 5-hmCyt, 5-fCyt, 5-caCyt, and 5-hmUra, we used a haploid HAP1 line, in which the producer had made a functional knockout of the TET1, TET2 or TET3 gene. These cells were a model of a disease in which the activity of one of the mentioned proteins is lost. Moreover, both wild-type and with single (SKO) functional knockouts in the genes TET1, TET2, and TET3 were cultured for 24 hours in the presence of 100 µmol/l of vitamin C to answer the question if vitamin C supplementation can restore lost enzymatic activity due to functional knockout. The results of the experiment are presented in Figure 1. It is worth noting that regardless of the mutation status, we observed a significant increase in 5-hmCyt, 5-fCyt, 5-caCyt, and 5-hmUra compared to levels of 5-hmCyt, 5-fCyt, 5-caCyt and 5-hmUra observed in control cells cultured without vitamin C, despite characteristic differences in their levels between cells with individual TET protein knockouts. Moreover, the lowest levels of 5-hmCyt, and 5-fCyt, compared to Parental HAP1 cells, were observed in the functional knockout of TET2, while the lowest level of 5-caCyt was observed in cells with functional knockout of TET3. However, only in case of 5-hmCyt difference was statistically significant (p<0.001, two-way ANOVA test with Tukey’s HSD test, the results of all statistical analyses for this experiment are presented in the Supplementary Table 1 in the Additional File 2). We have also analyzed results as a fold change relative to the level observed in control cells – data presented in Figure 2 (and as log_2_ fold change in Supplementary Figure 4 in Additional File 2). In this context, vitamin C supplementation, although it induced an absolute increase in the levels of 5-hmCyt, 5-fCyt, 5-caCyt, and 5-hmUra, did not seem to change their proportions relative to the control cells and did not lead to a change in a specific, established profile of TET protein activity resulting from the introduced knockouts. Moreover, after such normalization, we observed significant decrease in the level of 5-hmCyt and 5-fCyt in HAP1 TET1KO (45% and 29%, respectively), HAP1 TET2KO (70% and 51%, respectively) and HAP1 TET3KO (21% and 30%, respectively) in comparison to Parental HAP1 cells. The results of all statistical analyses for data shown as a fold change are presented in the Supplementary Table 2 in the Additional File 2

**Figure 1.**
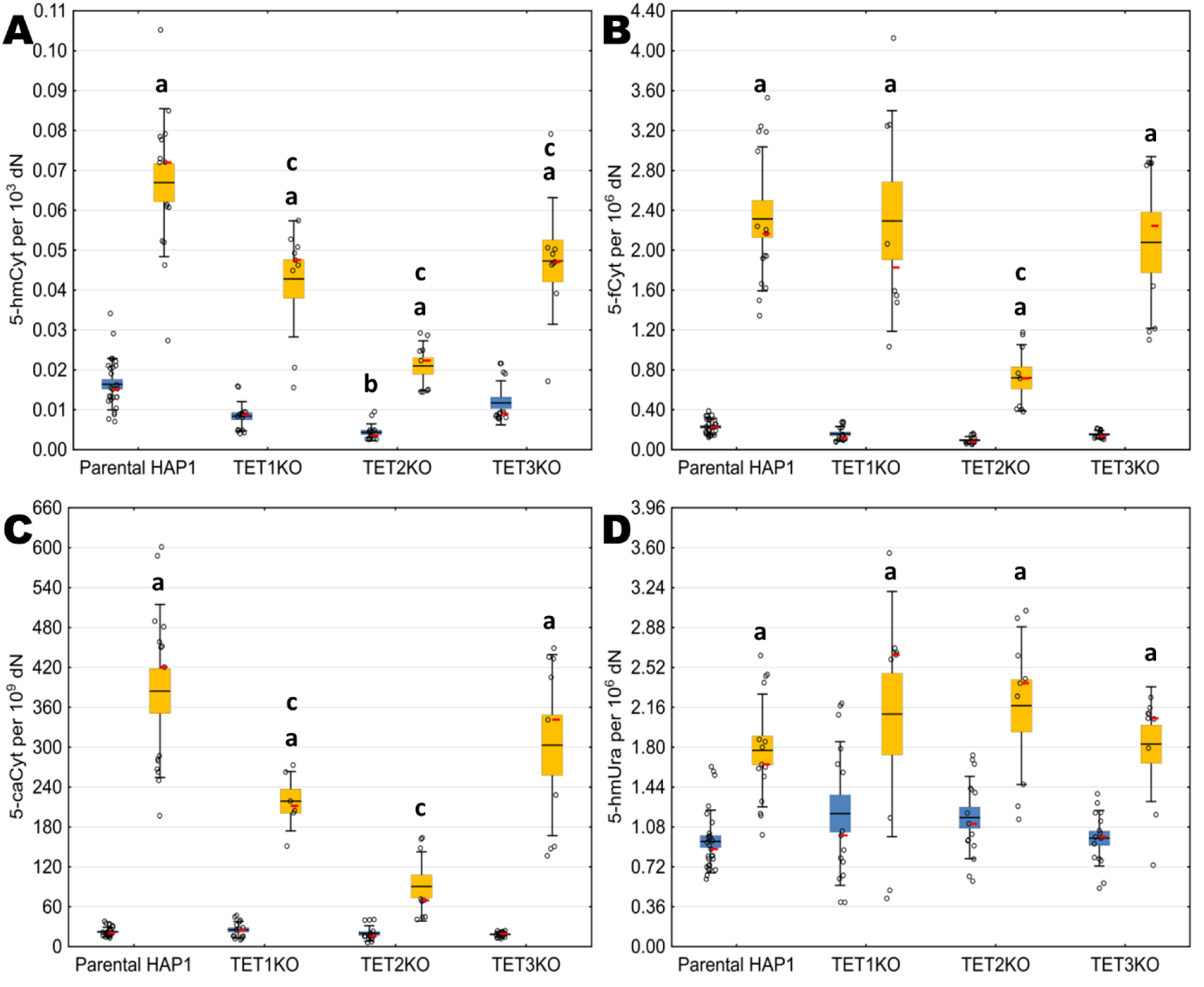
Content of modified nucleobases in HAP1 SKO cells after exposition to vitamin C. Content of modified nucleobases in DNA isolated form HAP1 cells (parental and with single TET knock-outs) cultured for 24 hours without vitamin C (blue bars) and with vitamin C in concentration of 100 µmol/l (yellow bars), presented as their individual level (A, B, C, D). A: 5-hmCyt; B: 5-fCyt; C: 5-caCyt, D: 5-hmUra. Box: mean ± standard error, whiskers: mean ± standard deviation, dots: raw data, red dash: median. Small letter “a” indicates significance between cells that were treated with vitamin C and untreated ones within cell line variant; small letter “b”: indicates significance in comparison to untreated Parental HAP1; small letter “c”: significance in comparison to Parental HAP1 exposed to vitamin C. All datasets are available in articlès additional file 1.

**Figure 2.**
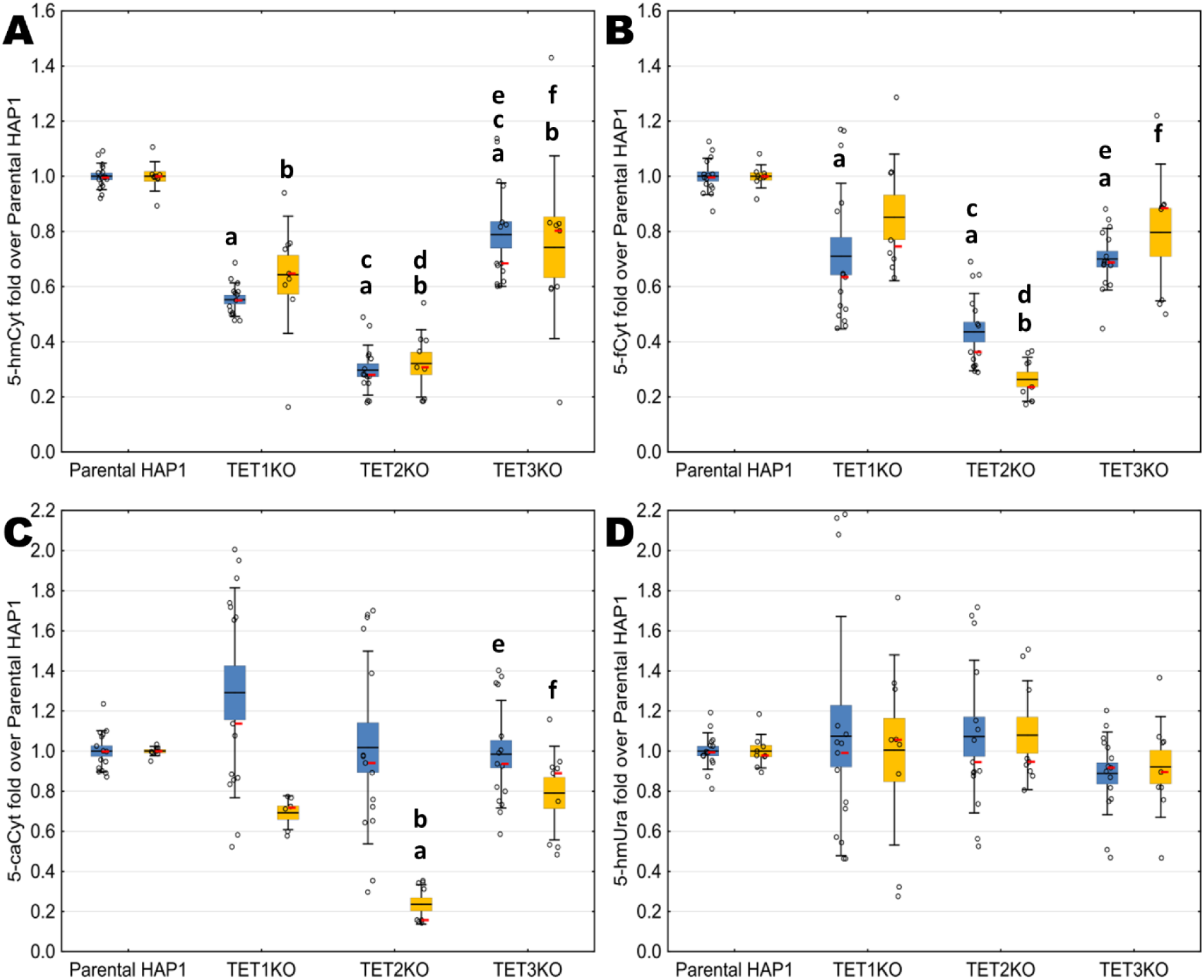
Content of modified nucleobases in HAP1 SKO cells represented as a fold change. Content of modified nucleobases in DNA isolated form HAP1 cells (parental and with single TET knock-outs) cultured for 24 hours without vitamin C (blue bars) and with vitamin C in concentration of 100 µmol/l (yellow bars), presented as their fold over level observed in parental line (A, B, C, D). A: 5-hmCyt; B: 5-fCyt; C: 5-caCyt, D: 5-hmUra. Box: mean ± standard error, whiskers: mean ± standard deviation, dots: raw data, red dash: median, small letter “a” indicates significance in comparison to untreated Parental HAP1, small letter “b” indicates significance in comparison to Parental HAP1 treated with vitamin C; small letters “c” and “d” indicate significance in comparison to respectively untreated and treated HAP1 TET1KO; small letters “e” and “f” indicate significance in comparison to respectively untreated and treated HAP1 TET2KO. All datasets are available in articlès additional file 1.

Moreover, to answer the question of whether the observed changes in the levels of TET protein activity products in individual knockouts could be explained by compensatory changes in the expression of the remaining TET proteins, we performed RT-qPCR analyses of individual *TET*s mRNA levels in SKO lines (see Supplementary Figure 5 in Additional File 2). However, these analyses did not reveal significant changes in the *TET* mRNA expression level in cells with single functional knock-outs that could indicate, that the observations from the chromatographic data are significantly altered by compensatory changes in the expression of the remaining TET proteins.

### Effect of ascorbic acid supplementation in HAP1 cell line with double knock-outs of TET genes

In order to determine the participation of individual TET family proteins in the generation of modified nucleobases of epigenetic importance, we decided to use HAP1 cells with functional knockout of two of the three *TET* genes, obtaining cell model in which only one TET protein remains active. Then, these cells were incubated for 24 hours in the presence of 100 µmol/l ascorbic acid, which allows for the estimation of the effect of this compound on the enzymatic activity of each member of the TET protein family. The results of the experiment are presented in Figure 3. It is worth noting that, similarly to the case of individual knockouts, supplementation of all cell lines with vitamin C caused a significant increase in the levels of 5-hmCyt, 5-fCyt, 5-caCyt, and 5-hmUra in comparison to levels of 5-hmCyt, 5-fCyt, 5-caCyt, and 5-hmUra observed in cells cultured without vitamin C. Moreover, observed level of 5-hmCyt in HAP1 TET1KO/TET2KO, HAP1 TET1KO/TET3KO and TET2KO/TET3KO was significantly decreased in comparison to level observed in Parental HAP1 cell line. The results of all statistical analyses for this experiment are presented in the Supplementary Table 3 in the Additional File 2.

**Figure 3.**
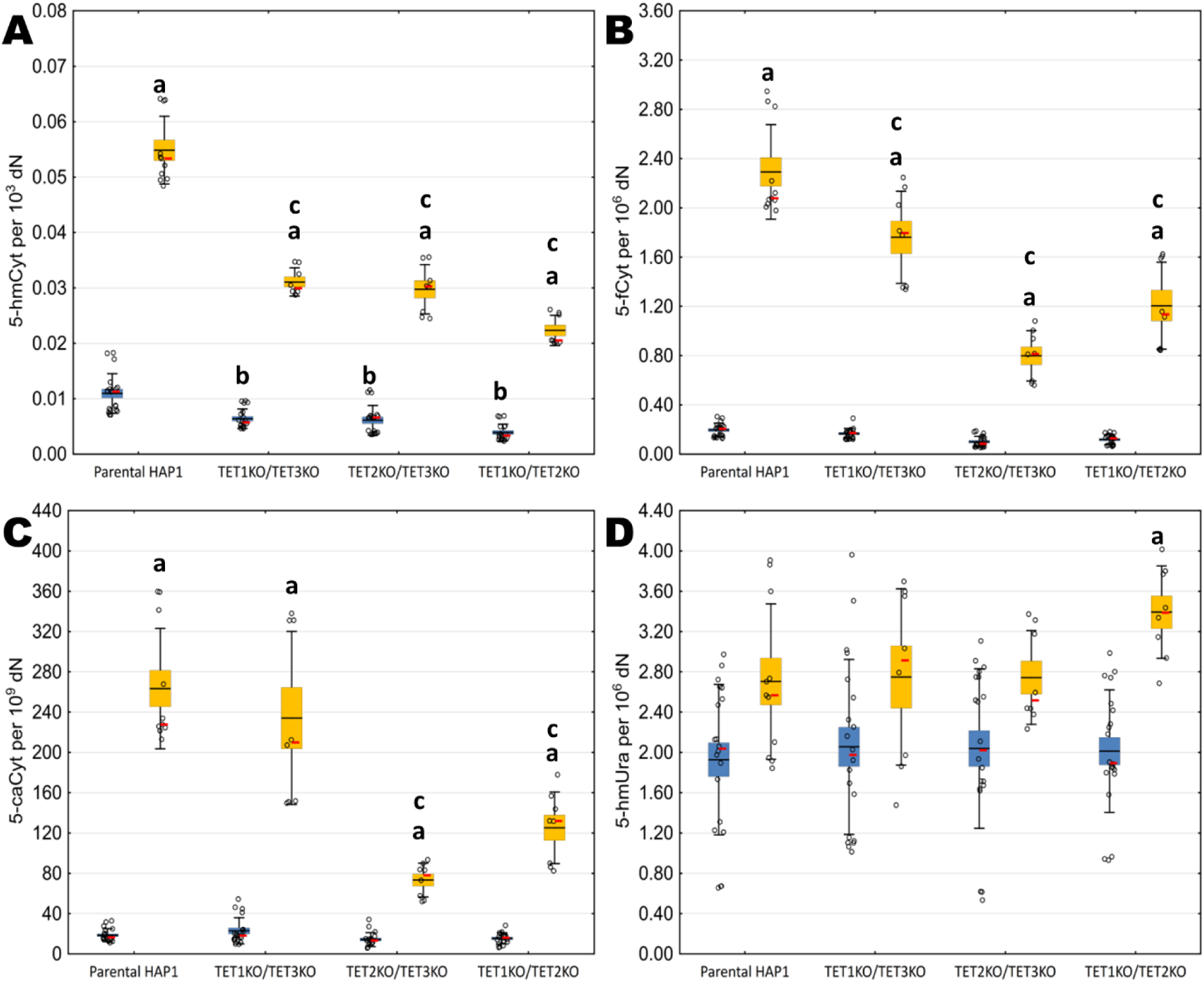
Content of modified nucleobases in HAP1 DKO cells after exposition to vitamin C. **Supplementary Figure 3 legend:** Content of modified nucleobases in DNA isolated form HAP1 cells (parental and with double TET knock-outs) cultured for 24 hours without vitamin C (blue bars) and with vitamin C in concentration of 100 µmol/l (yellow bars), presented as their individual level (A, B, C, D). A: 5-hmCyt; B: 5-fCyt; C: 5-caCyt, D: 5-hmUra. Box: mean ± standard error, whiskers: mean ± standard deviation, dots: raw data, red dot: median. All datasets are available in articlès additional file. Small letter “a” indicates significance between cells that were treated with vitamin C and untreated ones within cell line variant; small letter “b” indicates significance in comparison to untreated Parental HAP1; small letter “c” indicates significance in comparison to Parental HAP1 exposed to vitamin C. All datasets are available in articlès additional file 1.

We also decided to compare the results as a fold change relative to the level observed in control cells (presented in Figure 4 and as a log_2_ fold change in Supplementary Figure 6 in Additional File 2). In DKO, vitamin C supplementation, despite inducing an absolute increase in the levels of 5-hmCyt, 5-fCyt, 5-caCyt, and 5-hmUra did not changed the proportions of these modifications in relation to the levels observed in Parental HAP1 cells. Moreover, when data were presented as a fold change, we have observed significant decrease in the level of 5-hmCyt and 5-fCyt in all DKO cell lines (TET1KO/TET3KO: 40% and 13%, respectively; TET2KO/TET3KO: 46% and 50%, respectively and TET1KO/TET2KO: 65% and 41%, respectively) in comparison to Parental HAP1. The results of all statistical analyses for data shown as a fold change are presented in the Supplementary Table 4 in the Additional File 2

**Figure 4.**
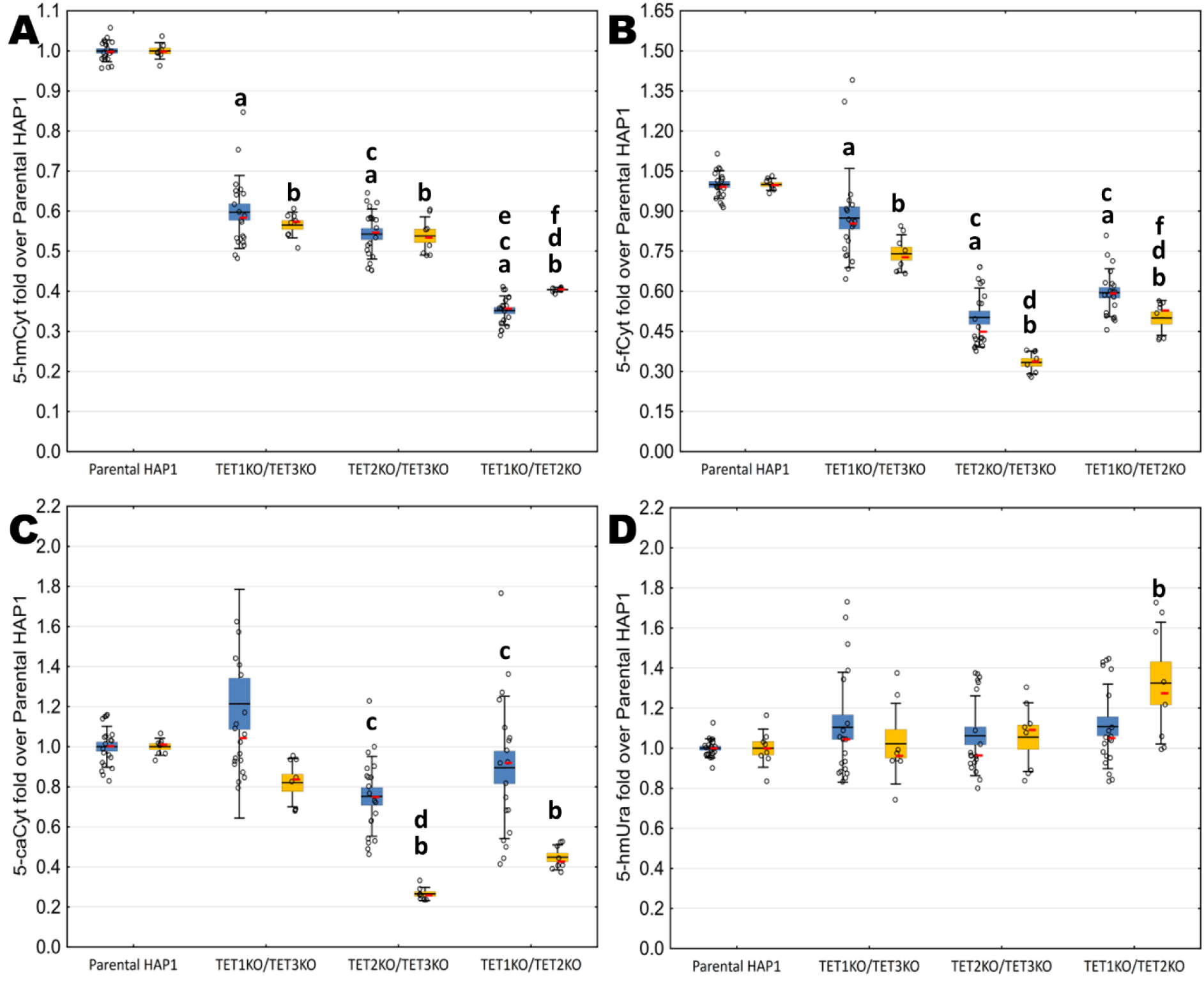
Content of modified nucleobases in HAP1 DKO cells represented as a fold change. Content of modified nucleobases in DNA isolated form HAP1 cells (parental and with double TET knock-outs) cultured for 24 hours without vitamin C (blue bars) and with vitamin C in concentration of 100 µmol/l (yellow bars), presented as their fold over level observed in parental line (A, B, D, F). A: 5-hmCyt; B: 5-fCyt; C: 5-caCyt, D: 5-hmUra. Box: mean ± standard error, whiskers: mean ± standard deviation, dots: raw data, red dash: median, small letter “a” indicates significance in comparison to untreated Parental HAP1, small letter “b” indicates significance in comparison to Parental HAP1 treated with vitamin C; small letters “c” and “d” indicate significance in comparison to respectively untreated and treated HAP1 TET1KO/TET3KO; small letters “e” and “f” indicate significance in comparison to respectively untreated and treated HAP1 TET2KO/TET3KO. All datasets are available in articlès additional file 1.

### Effect of ascorbic acid supplementation in HAP1 TET1KO/TET3KO cell line with reduced TET2 expression

In order to create a cell model, which was intended to simulate the triple knockout (TKO) effect, we used HAP1 TET1KO/TET3KO line, in which we silenced the expression of the TET2 gene using anti-TET2 siRNA molecules. Then, those near-TKO state cells, as well as, control cells treated with scrambled siRNA were exposed to 100 µmol/l of vitamin C for 24 hours. Results of this experiment are shown in Figure 5 (and as box-and-whisker plots in Supplementary Figure 7 in Additional File 2). We have not observed any significant differences (p>0.05) in the levels of 5-hmCyt, 5-fCyt, 5-caCyt and 5-hmUra between control cells and cells transfected with scrambled siRNA. Moreover, as expected, exposition to 100 µmol/l vitamin C significantly increased genomic content of 5-hmCyt, 5-fCyt, 5-caCyt and 5-hmUra in HAP1 TET1KO/TET3KO cell line. We have also observed significant decrease of 5-hmCyt and 5-fCyt, but not 5-caCyt and 5-hmUra levels in cells transfected with antiTET2 siRNA in comparison with control cells. Moreover, in cells transfected with antiTET2 siRNA, exposed to 100 µmol/l of vitamin C, we have observed significant increase in genomic content of 5-hmCyt, 5-fCyt, 5-caCyt and 5-hmUra in comparison to cells transfected with antiTET2 siRNA without exposition to vitamin C. We have also observed significant increase in genomic content of 5-hmUra in cells transfected with antiTET2 siRNA, exposed to 100 µmol/l of vitamin C in comparison to control cells. In addition, we have also observed significant increase in 5-hmCyt, 5-fCyt and 5-caCyt levels in control cells transfected with scrambled siRNA and exposed to 100 µmol/l of vitamin C when compared to cells exposed only to 100 µmol/l of vitamin C. The results of all statistical analyses for this experiment are presented in the Supplementary Table 5 in the Additional File 2.

**Figure 5.**
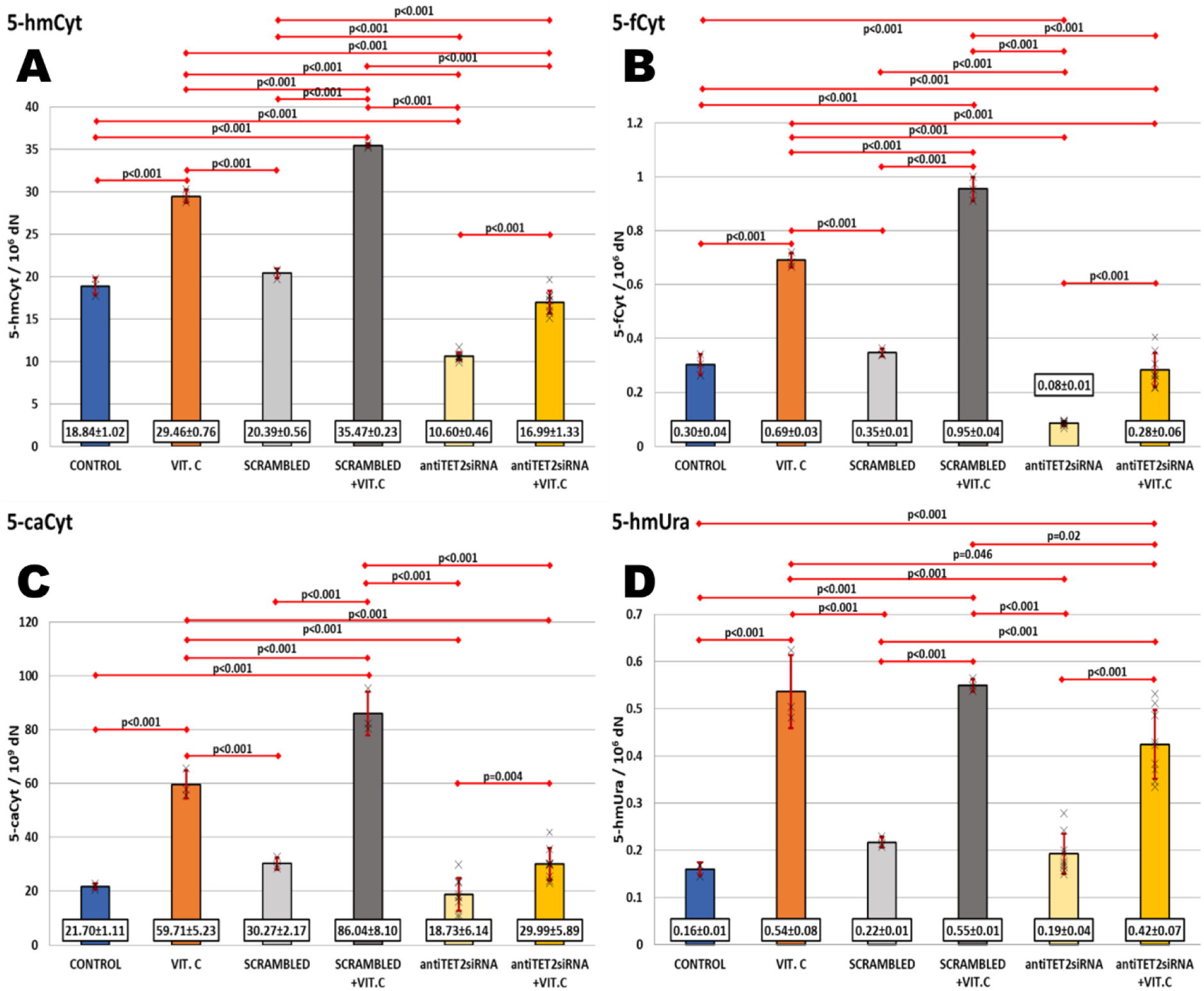
Content of modified nucleobases in HAP1 near-TKO state cells after exposition to vitamin C. Content of modified nucleobases in DNA isolated form HAP1 TET1KO/TET3KO cell line after 24 hours of transfection with antiTET2 siRNA duplexes and exposition to 100 µmol/l of vitamin C. A: 5-hmCyt; B: 5-fCyt; C: 5-caCyt, D: 5-hmUra. Data presented as mean ± standard deviation, x: raw data. All datasets are available in articlès additional file 1.

## DISCUSSION

Mammalian *TETs* (1–3), in addition to actively participating in the active demethylation process, may also individually deposit 5-hmCyt, 5-fCyt, 5-caCyt (possibly also 5-hmUra), and, in this way, may add new layers of epigenetic control in transcriptional regulation, cellular development, and lineage specification (reviewed in (50).

Loss-of-function mutations in *TET2*, which resemble the KO state, have been described in various hematological malignancies, including 7% to 10% of acute myeloid leukemia patients (discussed in (51)). Pronier et al. demonstrated that *TET2* knockdown skews human progenitor differentiation toward the myeloid lineage (30). Although *TET1* and *TET3* mutations are infrequent in hematologic malignancies, recent studies indicate that *TET1* also retains a crucial regulatory role in the oncogenic transformation of hematopoietic cells (52). Thus, using *TET* KO myeloid leukemia cells (HAP1), we ask the question of whether different paralogs of *TETs* have distinct, separate contributions to generating active demethylation products (the whole spectrum) in the malignant cells.

Regulation of the enzymatic activity of TET proteins may precisely control the patterns of 5-mCyt and its oxidized derivatives using vitamin C, which enhances TET activity and significantly increases levels of 5-hmCyt, 5-fCyt,5-caCyt, and 5-hmUra. Therefore, we also seek to determine whether vitamin C can compensate for the lack of a particular paralog of *TET* in KO cells.

It has been demonstrated that genomic levels of 5-hmCyt are profoundly decreased in many types of human malignancies, and the levels are distinctive epigenetic indicators of clinical outcomes and correlate with an increased risk of neoplastic progression (53). A decrease in 5-hmCyt levels may be due to mutations of *TETs*. *TET2* is the most frequently mutated gene in myeloid neoplasia. Thus, *TET2* loss of function alone is involved in leukemic progression or aneuploidy (54, 55).

Our study used our recently developed, highly sensitive, and specific methodology to determine how individual TETs shape a pattern of active demethylation products in HAP1 cells with single and double functional knockouts in TET genes.

We found the essential role of TET2 in all the steps of iterative oxidation and the significant involvement of TET1 in the formation of higher-order oxidation products, namely 5-hmCyt and 5-fCyt. Our methodology also enables us to reliably and quantitatively analyze the 5-caCyt level in genomic DNA, and we found a similar involvement of TET paralogs in its formation to that characteristic for a 5-fCyt generation.

Vitamin C indirectly enhances the activity of most α-KG/Fe^2+^ - dependent dioxygenases, including *TET* paralogues, by the most accepted mechanism that vitamin C efficiently recharges the active site Fe^3+^ to Fe^2+^ (56). Then, the active site Fe^2+^ cleaves the O‒O bond to produce succinate, CO_2_, and hydroxylated product. Vitamin C may thus increase the throughput of intact TETs, i.e., it may restore hydroxymethylation and thus reverse the epigenetic phenotype of the malignant cells.

All three TET proteins share a high degree of homology within their catalytic domains (57, 58); vitamin C may thus improve the function of intact TET2 or increase the compensatory enzymatic activity of TET1/TET3 in cases with TET2 inactivation. Such effects may restore or improve hydroxymethylation and potentially reverse the epigenetic consequences caused by *TET2* deficiency.

This hypothesis is supported by research conducted by Guan Y. et al., in which it was found that vitamin C directly binds the catalytic domain of TET2 in a dose-dependent manner, significantly enhancing its enzymatic activity. It facilitates the redox conversion of Fe^3+^ to Fe^2+^ at the catalytic site, which is essential for TET2 function (59). Additionally, it was found that vitamin C can act as an alternate oxygen acceptor, supporting TET2 activation under certain conditions. However, the response to vitamin C varied based on the specific TET2 mutations and regulatory factors influencing TET2 stability and activity. Moreover, vitamin C effects were observed in both heterozygous and biallelic TET2MT settings, as well as in Tet2-null cells, likely due to its ability to enhance the activity of other TET enzymes (TET1, TET3) and its broader cellular effects (45).

To test this theoretical assumption, we exposed Parental HAP 1 cells and HAP1 with individual and double knockouts to vitamin C (100 µmol/l). HAP1 cells, like most malignant tissues, have a profound decrease of 5-hmCyt. The addition of vitamin C to Parental HAP1 substantially increases higher-order oxidative products of TET activity and, in the case of TET1 and TET3, KO cells, to some degree, regenerate the level seen in Parental HAP1. However, in TET2 KO cells, the level increases only slightly, far below this characteristic for the parental line. In good agreement with our recently published work (60), we have found that ascorbate stimulated a moderate (2-to 3-fold) increase in the level of 5-hmCyt and a substantial increase in 5-fCyt and 5-caCyt levels. A similar picture was found in TET1KO and TET3KO cells. However, vitamin C increased the level of 5-hmCyt several times in TET2 KO cells compared with non-exposed cells, but the level is still lower than those observed in stimulated Parental HAP1 cells indicating incomplete compensation for TET2 loss. More pronounced increases were found in the case of 5-fCyt and 5-caCyt in all TETs KO.

Products of 5-mCyt demethylation, namely 5-hmCyt and 5-fCyt, dropped, respectively, to about 45% and 29% of their Parental levels in TET1 KO, to 70% and 51% in TET2 KO cells, and to 21% and 30%, in TET3KO. A similar decrease was also observed in TET1KO/TET2KO cells (65% and 41%, respectively).

Intriguingly, after activation of TETs with vitamin C to this set of cells, these values still stay roughly the same, which, in turn, strongly suggests that a lack of specific TET (1 or 2) cannot be fully compensated by the other enzyme(s). Although ascorbic acid strongly potentiates individual TET proteins, our data suggest that deletion of one of the TET enzymes, especially TET2 and, to a lesser degree, TET1, cannot be compensated for by the increased activity of the other TET family members. However, based on our results from siRNA experiments, it is still possible to enhance residual TET activity to the degree that overcame its activity impairment arising from a reduced expression, which constitutes the premise to supplement patients with disorders in the expression/activity of individual TET proteins. Moreover, to our surprise, our data showed that vitamin C strongly enhances the generation of 5-hmUra in HAP1 TET1KO/TET3KO cells with impaired TET2 activity, which suggests that TET activity is not the only or predominant source of this modification in human DNA. The observed increase in 5-hmUra levels in these cells may result from the compensatory action of alternative pathways for forming 5-hmUra, such as the intensification of DNA deamination processes, for example, through increased expression/increased activity of DNA deaminases. However, the literature has not shown that vitamin C directly affects their expression/activity. It also cannot be ruled out that part of the 5-hmUra pool observed in these cells is formed due to non-enzymatic thymine oxidation, such as ROS activity. Interestingly, recent work by Jin G. B. et al. (61) demonstrated that several mutations in the TET2 catalytic domain, like N1387T, result in substrate shift from 5-methylcytosine towards thymine and subsequently in elevated levels of genomic content of 5-hydroxymethyluracil. However, taking into consideration the fact that in this experiment and our previous work (60), vitamin C increased level of 5-hmUra in supplemented cells, we hypothesize that vitamin C, most probably by interacting with the TET catalytic domain (as shown in (45)) may act similarly to substrate-shifting mutations and enhance TET activity towards thymine. However, this hypothesis should be further investigated in future experiments.

Genetic studies strongly suggest that TET proteins are functionally nonredundant, most likely due to varying expression levels and recruitment mechanisms (62). Moreover, interaction with multiple binding partners may recruit or exclude TETs from specific genomic sites (50).

In agreement with this suggestion, our results pointed out that every single TET protein has a specific, conserved function, which the paralogs cannot take over. We hypothesize that in addition to the chemical distributive feature of TETs, the enzymes are also characterized by distributive locale in the genome, i.e., individual TET protein is localized in particular, likely tissue (set of genes) specific territory.

It is established that TET’s preferred substrate is 5-mCyt over 5-hmCyt (37). Thus, TET predominantly formed 5-hmCyt. Moreover, the level of this modification may be a good predictor of TET2 mutations. However, our recently published paper found that a good predictor/marker of TET2 mutations in patients with myelodysplastic syndromes was 5-fCyt and 5-caCyt (63). Similarly to the aforementioned results, we demonstrated several-fold decreased levels in TET2 KO HAP1 cells, not only for 5-hmCyt but also for 5-fCyt.

As mentioned above, loss of function mutations of TET genes, especially TET2, are prevalent in hematological malignancies. Moreover, the direct link between vitamin C and TET function in the progression of many hematological disorders was demonstrated (33–36). Cancer patients, including those with hematological malignancies, are vitamin C – deficient. Our results suggest that even in the case when TET2 is dysfunctional, vitamin C supplementation stimulation with ascorbic acid may be helpful to recover the normal demethylation process partially, improve the function of TETs, restore epigenetic phenotype, and potentially reverse the epigenetic consequences of the enzyme deficiency.

## CONCLUSIONS

Summing up, our results demonstrated that in HAP1 cells: i/ TETs have a distinct, separate contribution to generating active demethylation products (the whole spectrum), and the absence of individual TET paralog is linked with the specific pattern of active demethylation products of DNA; ii/ for the first time we have demonstrated that despite significant potentiation of TETs activity with ascorbic acid this pattern is preserved after vitamin C treatment, however, reinforced by a certain factor, what in turn suggest that deletion of one of the TET enzyme cannot be compensated for by increased activity of the other TET family members; iii/ TET2 is indispensable for vitamin C dependent regeneration of the deficiency of DNA active demethylation products in TET 1/3 KO myeloid leukemia cells.

## LIST OF ABBREVIATIONS

5-caCyt - 5-carboxycytosine

5-fCyt – 5-formylcytosine

5-hmCyt −5-hydroxymethylcytosine

5-hmUra - 5-hydroxymethyluracil

5-mCyt – 5-methylcytosine

DKO – double functional knockout of TET genes

SKO - single functional knock-outs of TET genes

TET - ten-eleven translocation family proteins

TKO – Total knock-out of TET genes

## DECLARATIONS

### Ethics approval and consent to participate

Not applicable

### Consent for publication

Not applicable

### Availability of data and materials

The dataset supporting the conclusions of this article is included within the article (and its additional file 1).

### Authors’ contributions

**M.G.** performed cell cultures and siRNA experiments, prepared samples for 2D-UPLC-MS/MS analyses, performed statistical analyses, prepared figures, and wrote the manuscript; **M.S.** performed DNA isolation, enzymatic digestion of nucleic acids, participated in 2D-UPLC-MS/MS assays, and manuscript preparation; **A.W.** performed cell cultures, and ascorbic acid supplementation experiments, participated in manuscript preparation; **T.D.** performed RT-qPCR analyses; **R.O.** participated in manuscript preparation, provided substantive supervision of the work, was responsible for the study concept; **D.G.** participated in 2D-UPLC-MS/MS analyses, in statistical analysis of data, and manuscript preparation. All authors read and approved the final version of the manuscript.

## Acknowledgements

All authors are members of the University Center of Excellence “Towards Personalized Medicine” operating under the Excellence initiative – Research University.

While preparing this work, the authors used Grammarly (v. 1.0.37.777) and ChatGPT-4o to improve language and readability. After using these tools, the authors reviewed and edited the content as needed and are taking full responsibility for the publication’s content.

