## Additional file 2 for "Loss of TET2 activity limits the ability of vitamin C to activate DNA demethylation in human HAP1 cells"

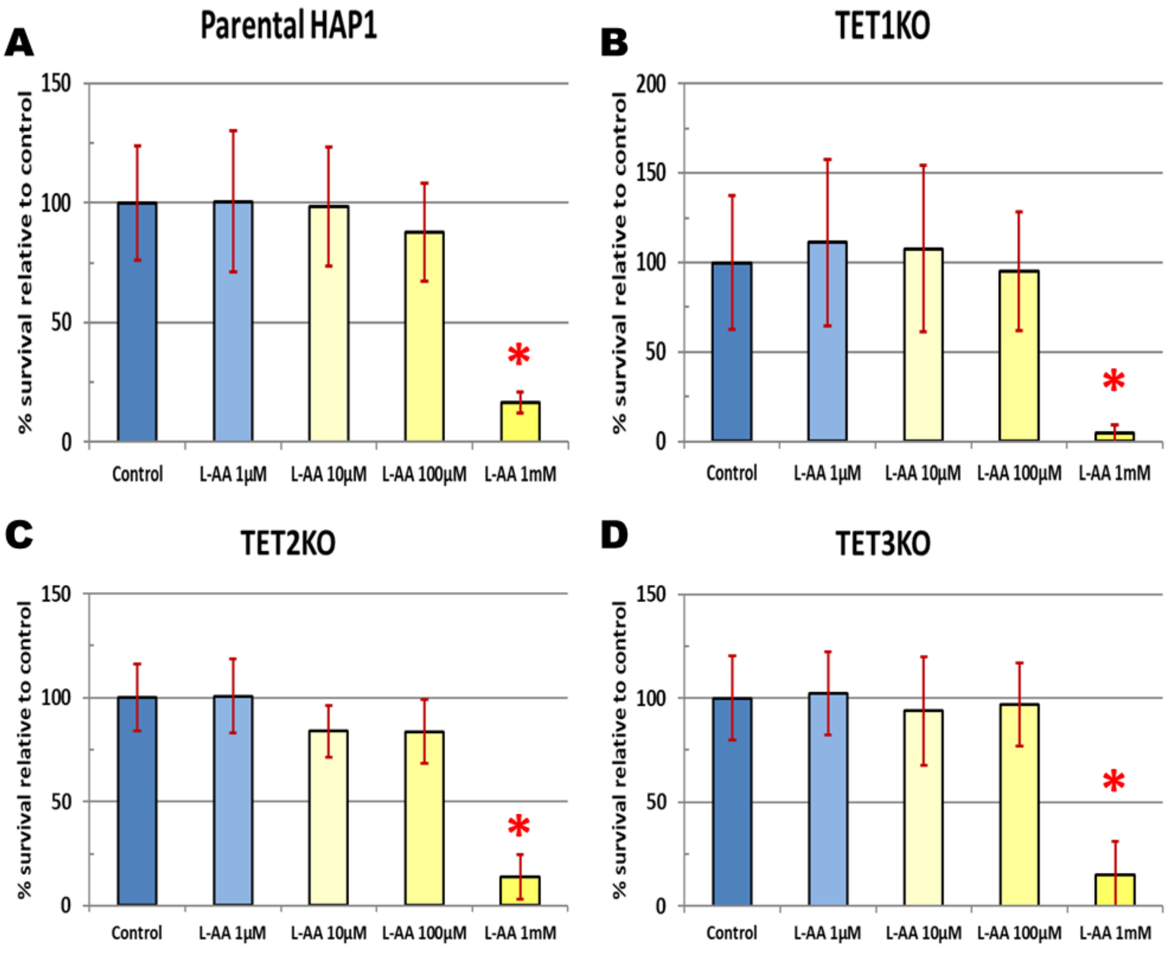


**Supplementary Figure 1 title:** Analysis of the survival of HAP1 cells exposed for 24h to various concentration of vitamin C.

**Supplementary Figure 1 legend:** Results of MTT assay are shown as % survival relative to control cells (mean ± standard deviation). L-AA – vitamin C, red asterisk indicates statistical significance.

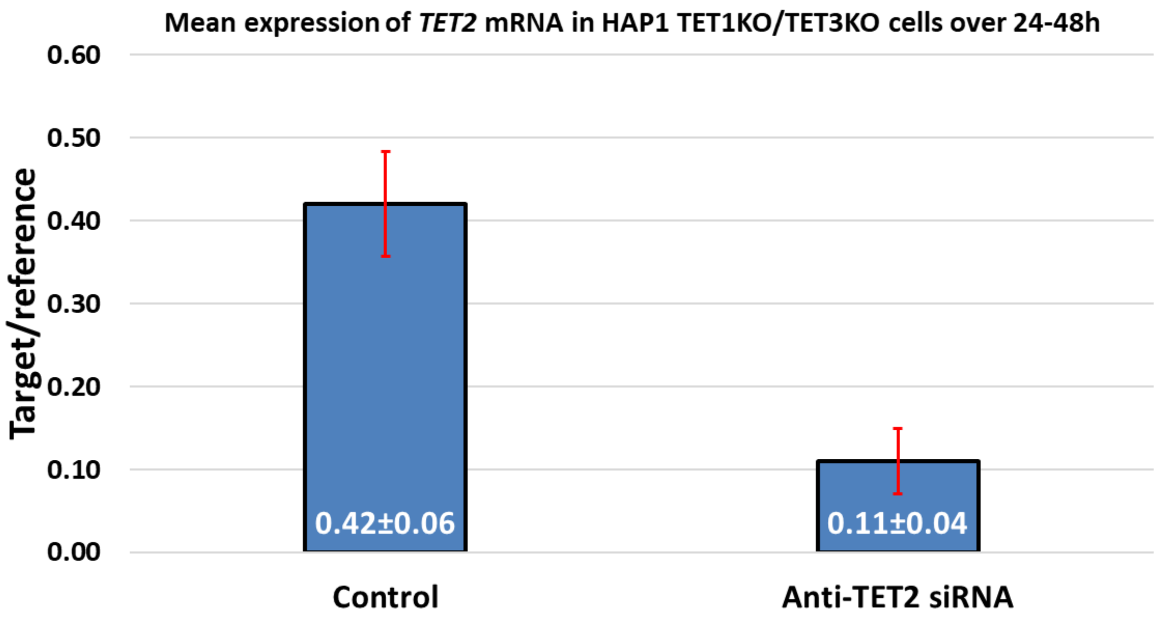


**Supplementary Figure 2 title**: Mean expression of *TET2* mRNA in HAP1 TET1KO/TET3KO cells after 24-48 hours of transfection.

**Supplementary Figure 2 legend:** Results are shown as mean ± standard deviation. *TET2* expression analysis was performed following the methodology described in: Starczak, M., Zarakowska, E., Modrzejewska, M. et al. In vivo evidence of ascorbate involvement in the generation of epigenetic DNA modifications in leukocytes from patients with colorectal carcinoma, benign adenoma and inflammatory bowel disease. J Transl Med 16, 204 (2018). https://doi.org/10.1186/s12967-018-1581-9

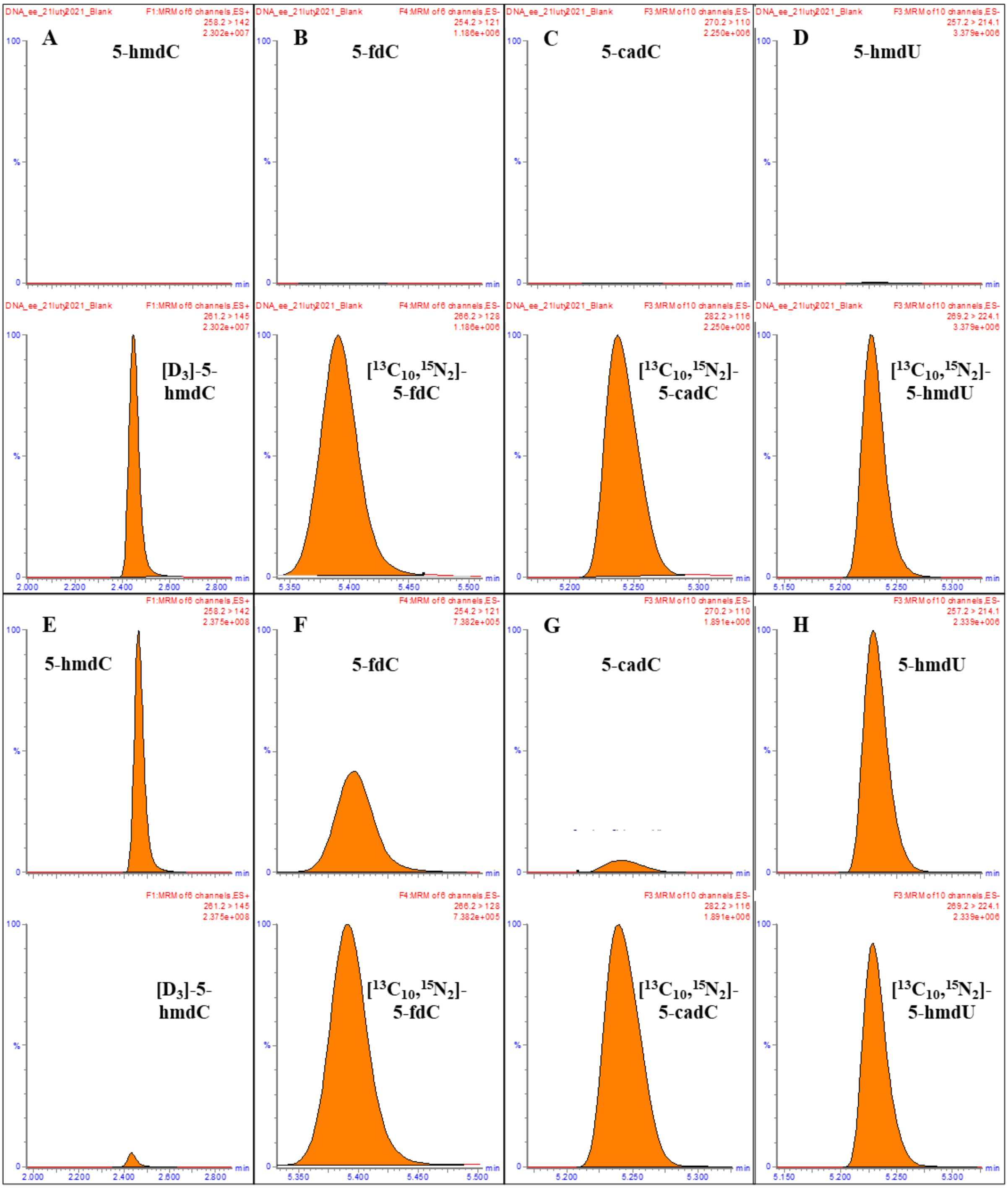


**Supplementary Figure 3 title:** Example chromatograms from 2D-UPLC-MS/MS analyses of the content of TET proteins activity products in HAP1 cell DNA.

**Supplementary figure 3 legend: A, B, C, D** – Blank samples chromatograms: 5-(hydroxymethyl)-2'-deoxycytidine (5-hmdC), 5-formyl-2'-deoxycytidine (5-fdC), 5-carboxy-2'-deoxycytidine (5-cadC), 5-(hydroxymethyl)-2'-deoxyuridine (5-hmdU) respectively, **E, F, G, H** – samples treated with 100 µmol/l vitamin C.

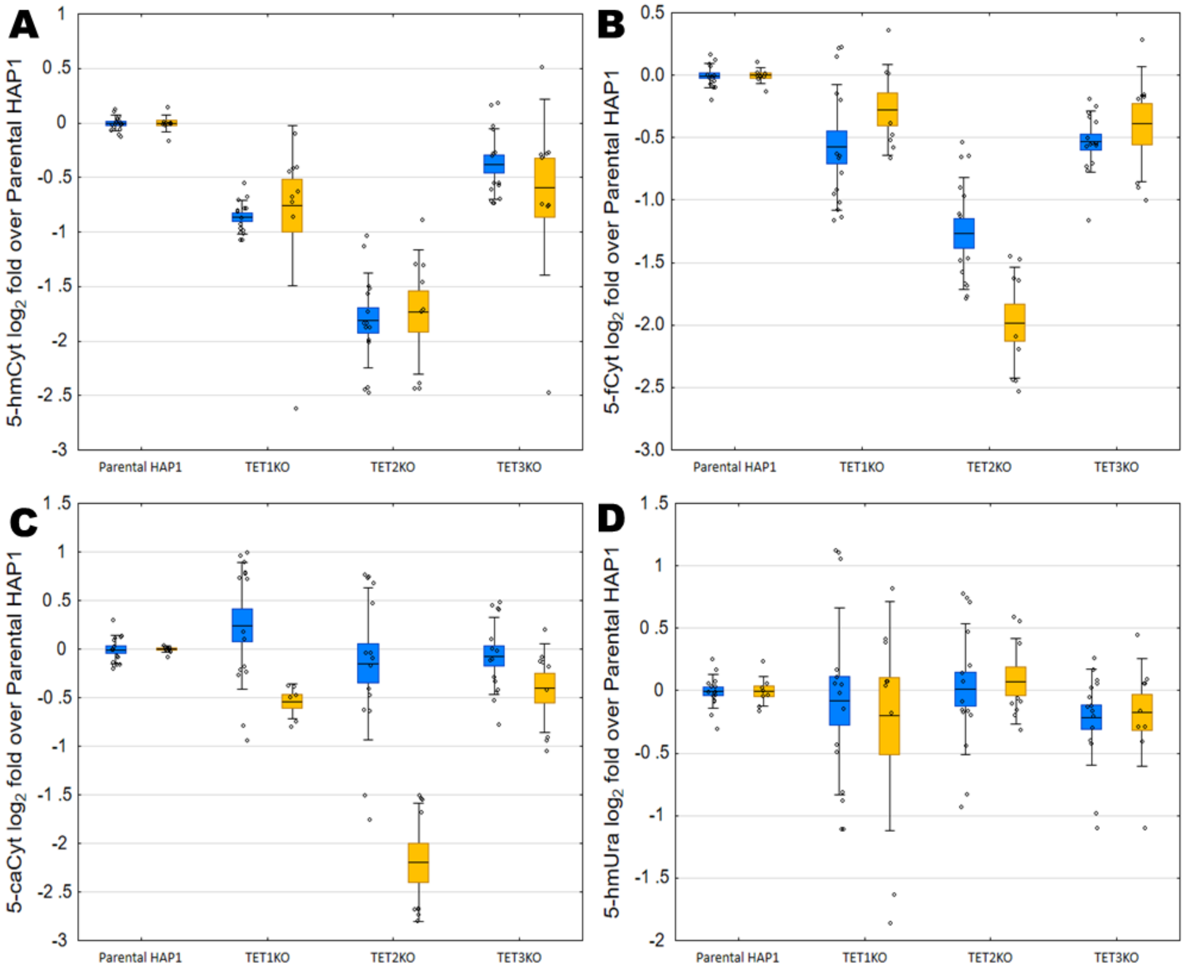


**Supplementary Figure 4 title:** Content of modified nucleobases in HAP1 SKO cells represented as a $\log_2$ fold change

**Supplementary Figure 4 legend:** Content of modified nucleobases in DNA isolated form HAP1 cells (parental and with single TET knock-outs) cultured for 24 hours without vitamin C (blue bars) and with vitamin C in concentration of 100 µmol/l (yellow bars), presented as their $\log_2$ fold over level observed in parental line (A, B, C, D). A: 5-hmCyt; B: 5-fCyt; C: 5-caCyt, D: 5-hmUra. Box: mean ± standard error, whiskers: mean ± standard deviation, dots: raw data.

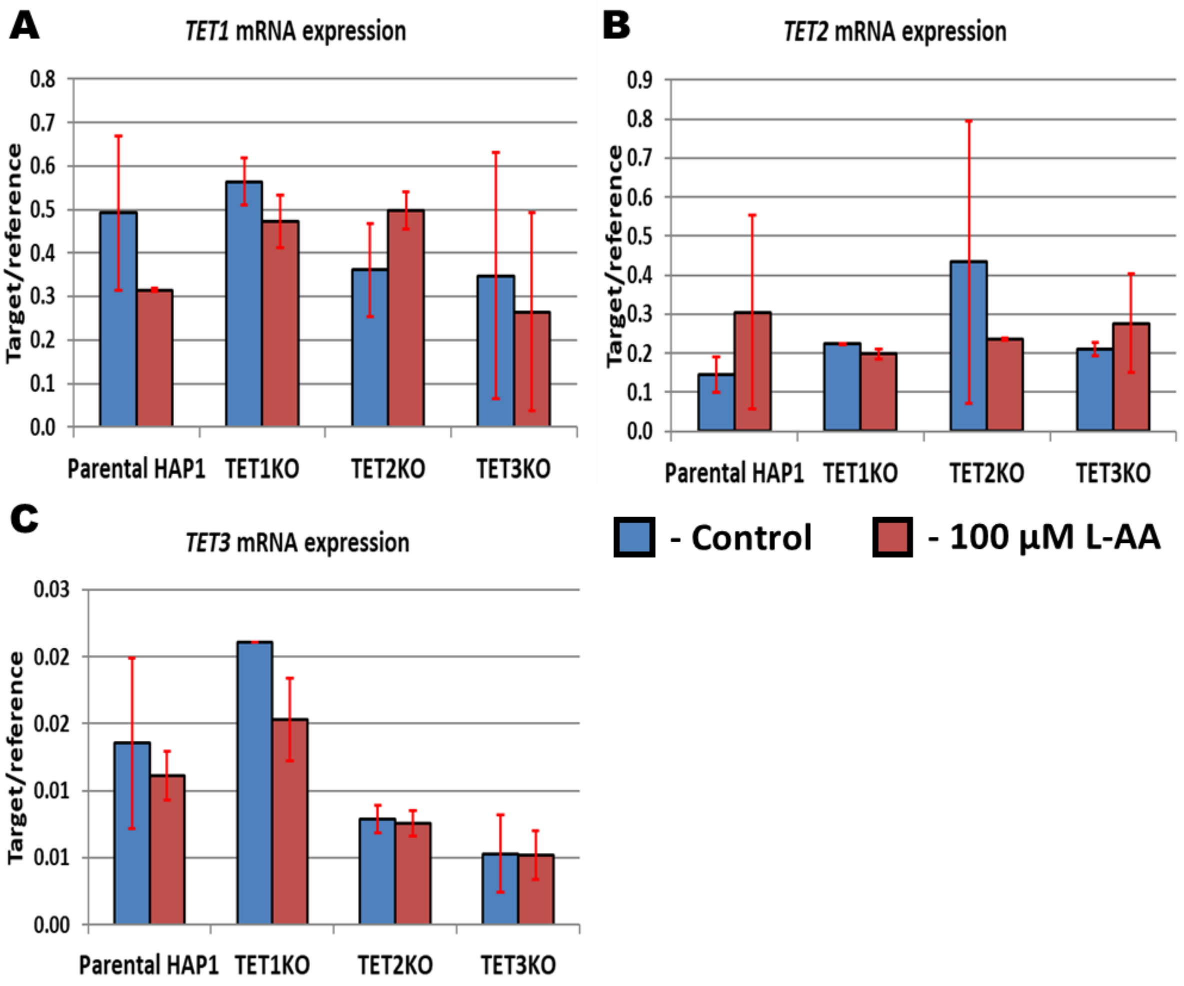


**Supplementary Figure 5 title:** Expression of *TET* genes mRNA in HAP1 cells
**Supplementary Figure 5 legend**: A: *TET1* mRNA, B: *TET2* mRNA, and C: *TET3* mRNA in HAP1 cells (Parental HAP1 and cells with functional knock-outs of individual *TET* genes). Blue bars represent control cells, and red bars represent cells after exposure to 100 µmol/l of vitamin C (L-AA). Data are shown as a mean ± standard deviation. Gene expression analysis was performed following the methodology described in: Starczak, M., Zarakowska, E., Modrzejewska, M. et al. In vivo evidence of ascorbate involvement in the generation of epigenetic DNA modifications in leukocytes from patients with colorectal carcinoma, benign adenoma and inflammatory bowel disease. J Transl Med 16, 204 (2018). https://doi.org/10.1186/s12967-018-1581-9.

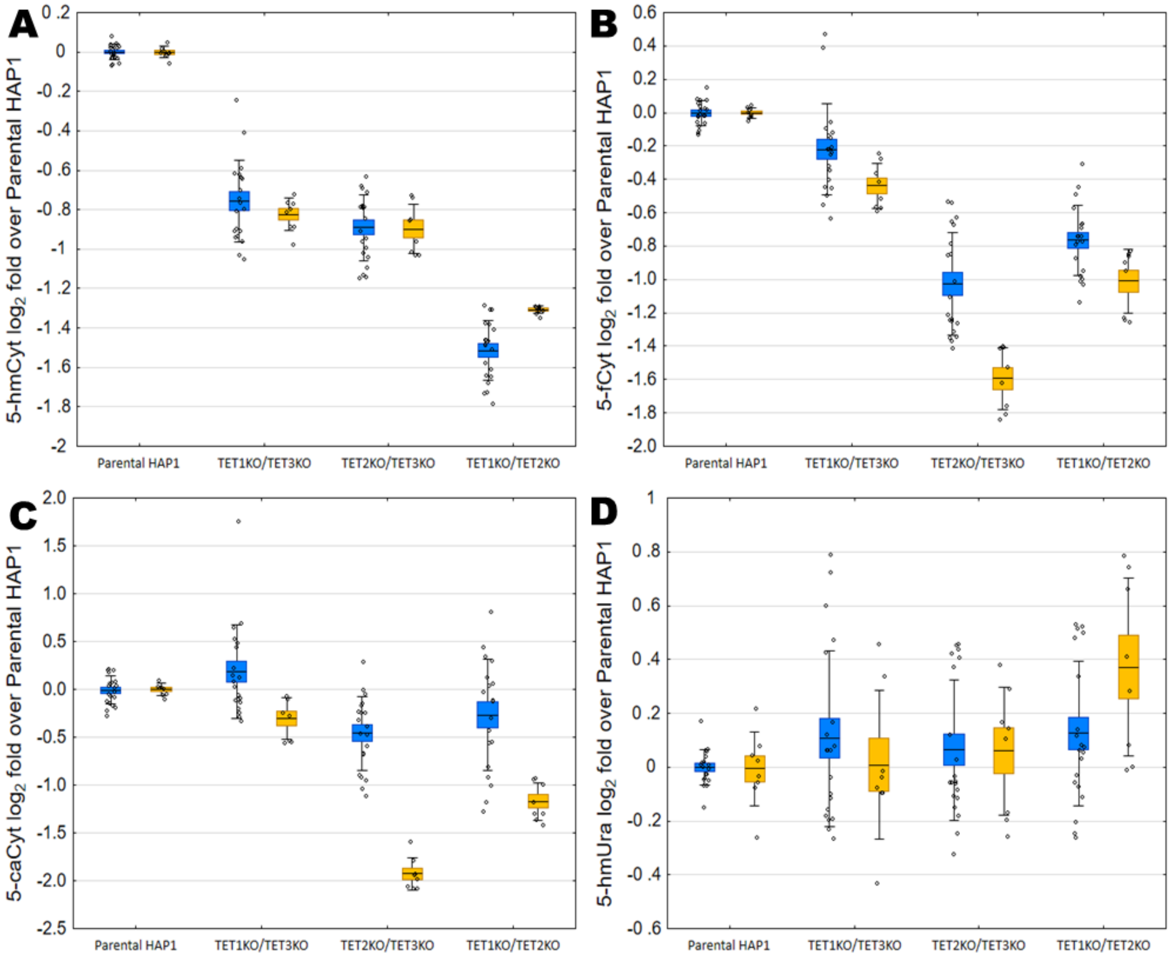


**Supplementary Figure 6 title:** Content of modified nucleobases in HAP1 DKO cells represented as a $\log_2$ fold change

**Supplementary Figure 6 legend:** Content of modified nucleobases in DNA isolated form HAP1 cells (parental and with double TET knock-outs) cultured for 24 hours without vitamin C (blue bars) and with vitamin C in concentration of 100 µmol/l (yellow bars), presented as their $\log_2$ fold over level observed in parental line (A, B, C, D). A: 5-hmCyt; B: 5-fCyt; C: 5-caCyt, D: 5-hmUra. Box: mean ± standard error, whiskers: mean ± standard deviation, dots: raw data.

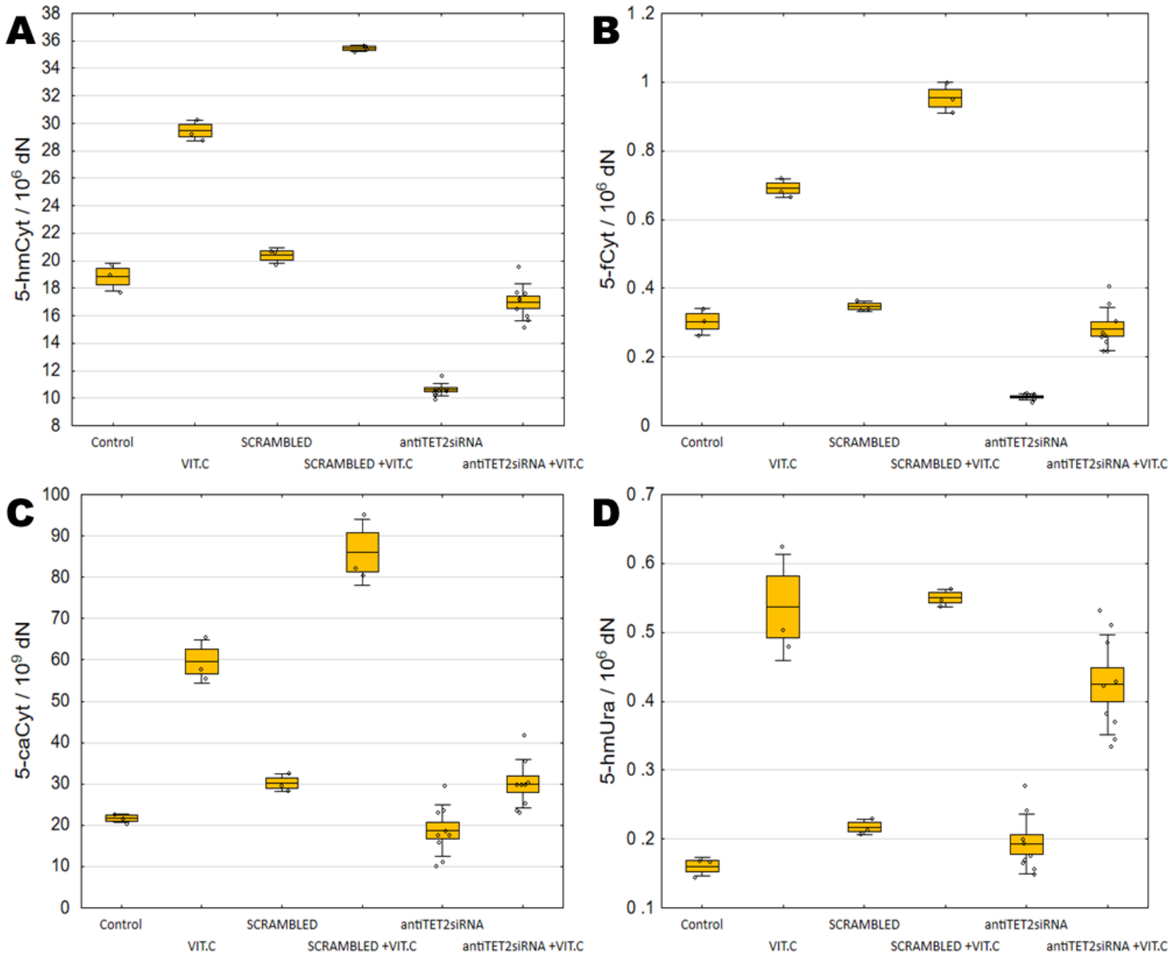


**Supplementary Figure 7 title:** Content of modified nucleobases in HAP1 near-TKO state cells after exposition to vitamin C

**Supplementary Figure 7 legend:** Content of modified nucleobases in DNA isolated form HAP1 TET1KO/TET3KO cell line after 24 hours of transfection with antiTET2 siRNA duplexes and exposition to 100 µmol/l of vitamin C. A: 5-hmCyt; B: 5-fCyt; C: 5-caCyt, D: 5-hmUra. Box: mean ± standard error, whiskers: mean ± standard deviation, dots: raw data.

| 5-hmCyt | Parental HAP1: Control | Parental HAP1: Vit. C | TET1KO: Control | TET1KO: Vit.C | TET2KO: Control | TET2KO: Vit. C | TET3KO: Control |
|---|---|---|---|---|---|---|---|
| TET3KO: Vit. C | 0.0001 | 0.0003 | 0.0001 | 0.9801 | 0.0001 | 0.0001 | 0.0001 |
| TET3KO: Control | 0.8285 | 0.0001 | 0.9841 | 0.0001 | 0.4745 | 0.3590 | |
| TET2KO: Vit. C | 0.9320 | 0.0001 | 0.0650 | 0.0004 | 0.0035 | | |
| TET2KO: Control | 0.0070 | 0.0001 | 0.9548 | 0.0001 | | | |
| TET1KO: Vit.C | 0.0001 | 0.0001 | 0.0001 | | | | |
| TET1KO: Control | 0.2096 | 0.0001 | | | | | |
| Parental HAP1: Vit. C | 0.0001 | | | | | | |

| 5-fCyt | Parental HAP1: Control | Parental HAP1: Vit. C | TET1KO: Control | TET1KO: Vit.C | TET2KO: Control | TET2KO: Vit. C | TET3KO: Control |
|---|---|---|---|---|---|---|---|
| TET3KO: Vit. C | 0.0001 | 0.9408 | 0.0001 | 0.9823 | 0.0001 | 0.0001 | 0.0001 |
| TET3KO: Control | 0.9996 | 0.0001 | 1.0000 | 0.0001 | 1.0000 | 0.0819 | |
| TET2KO: Vit. C | 0.1175 | 0.0001 | 0.0877 | 0.0001 | 0.0367 | | |
| TET2KO: Control | 0.9854 | 0.0001 | 0.9999 | 0.0001 | | | |
| TET1KO: Vit.C | 0.0001 | 1.0000 | 0.0001 | | | | |
| TET1KO: Control | 0.9998 | 0.0001 | | | | | |
| Parental HAP1: Vit. C | 0.0001 | | | | | | |

| 5-caCyt | Parental HAP1: Control | Parental HAP1: Vit. C | TET1KO: Control | TET1KO: Vit.C | TET2KO: Control | TET2KO: Vit. C | TET3KO: Control |
|---|---|---|---|---|---|---|---|
| TET3KO: Vit. C | 0.0001 | 0.0627 | 0.0001 | 0.2084 | 0.0001 | 0.0001 | 0.0001 |
| TET3KO: Control | 1.0000 | 0.0001 | 1.0000 | 0.0001 | 1.0000 | 0.1468 | |
| TET2KO: Vit. C | 0.1165 | 0.0001 | 0.2470 | 0.0059 | 0.1632 | | |
| TET2KO: Control | 1.0000 | 0.0001 | 1.0000 | 0.0001 | | | |
| TET1KO: Vit.C | 0.0001 | 0.0001 | 0.0001 | | | | |
| TET1KO: Control | 1.0000 | 0.0001 | | | | | |
| Parental HAP1: Vit. C | 0.0001 | | | | | | |

| 5-hmUra | Parental HAP1: Control | Parental HAP1: Vit. C | TET1KO: Control | TET1KO: Vit.C | TET2KO: Control | TET2KO: Vit. C | TET3KO: Control |
|---|---|---|---|---|---|---|---|
| TET3KO: Vit. C | 0.0012 | 1.0000 | 0.1130 | 0.9603 | 0.0764 | 0.8667 | 0.0066 |
| TET3KO: Control | 1.0000 | 0.0025 | 0.9465 | 0.0002 | 0.9795 | 0.0001 | |
| TET2KO: Vit. C | 0.0001 | 0.6285 | 0.0010 | 1.0000 | 0.0006 | | |
| TET2KO: Control | 0.9123 | 0.0498 | 1.0000 | 0.0019 | | | |
| TET1KO: Vit.C | 0.0001 | 0.8282 | 0.0032 | | | | |
| TET1KO: Control | 0.8238 | 0.0807 | | | | | |
| Parental HAP1: Vit. C | 0.0003 | | | | | | |

Supplementary Table 1. Tukey's HSD post hoc test results for comparisons of 5-hmCyt, 5-fCyt, 5-caCyt, and 5-hmUra levels in the DNA of HAP1 cells (Parental and with single functional knock-outs of *TET* genes) exposed for 24 hours to 100 µmol/l vitamin C.

| 5-hmCyt fold change | Parental HAP1: Control | Parental HAP1: Vit. C | TET1KO: Control | TET1KO: Vit.C | TET2KO: Control | TET2KO: Vit. C | TET3KO: Control |
|---|---|---|---|---|---|---|---|
| TET3KO: Vit. C | 0.0035 | 0.0137 | 0.0783 | 0.8649 | 0.0001 | 0.0001 | 0.9967 |
| TET3KO: Control | 0.0068 | 0.0318 | 0.0017 | 0.3352 | 0.0001 | 0.0001 | |
| TET2KO: Vit. C | 0.0001 | 0.0001 | 0.0129 | 0.0008 | 1.0000 | | |
| TET2KO: Control | 0.0001 | 0.0001 | 0.0005 | 0.0001 | | | |
| TET1KO: Vit.C | 0.0001 | 0.0002 | 0.8583 | | | | |
| TET1KO: Control | 0.0001 | 0.0001 | | | | | |
| Parental HAP1: Vit. C | 1.0000 | | | | | | |

| 5-fCyt fold change | Parental HAP1: Control | Parental HAP1: Vit. C | TET1KO: Control | TET1KO: Vit.C | TET2KO: Control | TET2KO: Vit. C | TET3KO: Control |
|---|---|---|---|---|---|---|---|
| TET3KO: Vit. C | 0.1066 | 0.1973 | 0.9338 | 0.9978 | 0.0002 | 0.0001 | 0.8811 |
| TET3KO: Control | 0.0002 | 0.0012 | 1.0000 | 0.4272 | 0.0009 | 0.0001 | |
| TET2KO: Vit. C | 0.0001 | 0.0001 | 0.0001 | 0.0001 | 0.2251 | | |
| TET2KO: Control | 0.0001 | 0.0001 | 0.0005 | 0.0001 | | | |
| TET1KO: Vit.C | 0.4532 | 0.5884 | 0.5274 | | | | |
| TET1KO: Control | 0.0003 | 0.0020 | | | | | |
| Parental HAP1: Vit. C | 1.0000 | | | | | | |

| 5-caCyt fold change | Parental HAP1: Control | Parental HAP1: Vit. C | TET1KO: Control | TET1KO: Vit.C | TET2KO: Control | TET2KO: Vit. C | TET3KO: Control |
|---|---|---|---|---|---|---|---|
| TET3KO: Vit. C | 0.7822 | 0.8640 | 0.0091 | 0.9991 | 0.7036 | 0.0098 | 0.8399 |
| TET3KO: Control | 1.0000 | 1.0000 | 0.1672 | 0.5658 | 1.0000 | 0.0001 | |
| TET2KO: Vit. C | 0.0001 | 0.0002 | 0.0001 | 0.1377 | 0.0001 | | |
| TET2KO: Control | 1.0000 | 1.0000 | 0.2903 | 0.4265 | | | |
| TET1KO: Vit.C | 0.5012 | 0.6120 | 0.0054 | | | | |
| TET1KO: Control | 0.2177 | 0.3930 | | | | | |
| Parental HAP1: Vit. C | 1.0000 | | | | | | |

| 5-hmUra fold change | Parental HAP1: Control | Parental HAP1: Vit. C | TET1KO: Control | TET1KO: Vit.C | TET2KO: Control | TET2KO: Vit. C | TET3KO: Control |
|---|---|---|---|---|---|---|---|
| TET3KO: Vit. C | 0.9995 | 0.9997 | 0.9656 | 0.9996 | 0.9684 | 0.9782 | 1.0000 |
| TET3KO: Control | 0.9880 | 0.9950 | 0.8266 | 0.9932 | 0.8359 | 0.8985 | |
| TET2KO: Vit. C | 0.9994 | 0.9997 | 1.0000 | 0.9998 | 1.0000 | | |
| TET2KO: Control | 0.9992 | 0.9997 | 1.0000 | 0.9998 | | | |
| TET1KO: Vit.C | 1.0000 | 1.0000 | 0.9998 | | | | |
| TET1KO: Control | 0.9990 | 0.9996 | | | | | |
| Parental HAP1: Vit. C | 1.0000 | | | | | | |

Supplementary Table 2. Tukey's HSD post hoc test results for comparisons of 5-hmCyt, 5-fCyt, 5-caCyt, and 5-hmUra fold change in the DNA of HAP1 cells (Parental and with single functional knock-outs of *TET* genes) exposed for 24 hours to 100 µmol/l vitamin C.

| 5-hmCyt | Parental HAP1: Control | Parental HAP1: Vit. C | TET1KO/ TET3KO: Control | TET1KO/ TET3KO: Vit.C | TET2KO/ TET3KO: Control | TET2KO/ TET3KO: Vit. C | TET1KO/ TET3KO: Control |
|---|---|---|---|---|---|---|---|
| TET1KO/TET3KO: Vit. C | 0.0001 | 0.0001 | 0.0001 | 0.0001 | 0.0001 | 0.0003 | 0.0001 |
| TET1KO/TET3KO: Control | 0.0001 | 0.0001 | 0.2153 | 0.0001 | 0.3545 | 0.0001 | |
| TET2KO/TET3KO: Vit. C | 0.0001 | 0.0001 | 0.0001 | 0.9902 | 0.0001 | | |
| TET2KO/TET3KO: Control | 0.0002 | 0.0001 | 1.0000 | 0.0001 | | | |
| TET1KO/TET3KO: Vit.C | 0.0001 | 0.0001 | 0.0001 | | | | |
| TET1KO/TET3KO: Control | 0.0005 | 0.0001 | | | | | |
| Parental HAP1: Vit. C | 0.0001 | | | | | | |

| 5-fCyt | Parental HAP1: Control | Parental HAP1: Vit. C | TET1KO/ TET3KO: Control | TET1KO/ TET3KO: Vit.C | TET2KO/ TET3KO: Control | TET2KO/ TET3KO: Vit. C | TET1KO/ TET3KO: Control |
|---|---|---|---|---|---|---|---|
| TET1KO/TET3KO: Vit. C | 0.0001 | 0.0001 | 0.0001 | 0.0001 | 0.0001 | 0.0009 | 0.0001 |
| TET1KO/TET3KO: Control | 0.8944 | 0.0001 | 0.9919 | 0.0001 | 1.0000 | 0.0001 | |
| TET2KO/TET3KO: Vit. C | 0.0001 | 0.0001 | 0.0001 | 0.0001 | 0.0001 | | |
| TET2KO/TET3KO: Control | 0.7531 | 0.0001 | 0.9549 | 0.0001 | | | |
| TET1KO/TET3KO: Vit.C | 0.0001 | 0.0001 | 0.0001 | | | | |
| TET1KO/TET3KO: Control | 0.9997 | 0.0001 | | | | | |
| Parental HAP1: Vit. C | 0.0001 | | | | | | |

| 5-caCyt | Parental HAP1: Control | Parental HAP1: Vit. C | TET1KO/ TET3KO: Control | TET1KO/ TET3KO: Vit.C | TET2KO/ TET3KO: Control | TET2KO/ TET3KO: Vit. C | TET1KO/ TET3KO: Control |
|---|---|---|---|---|---|---|---|
| TET1KO/TET3KO: Vit. C | 0.0001 | 0.0001 | 0.0001 | 0.0001 | 0.0001 | 0.0263 | 0.0001 |
| TET1KO/TET3KO: Control | 1.0000 | 0.0001 | 0.9952 | 0.0001 | 1.0000 | 0.0008 | |
| TET2KO/TET3KO: Vit. C | 0.0015 | 0.0001 | 0.0048 | 0.0001 | 0.0005 | | |
| TET2KO/TET3KO: Control | 0.9999 | 0.0001 | 0.9875 | 0.0001 | | | |
| TET1KO/TET3KO: Vit.C | 0.0001 | 0.4788 | 0.0001 | | | | |
| TET1KO/TET3KO: Control | 0.9999 | 0.0001 | | | | | |
| Parental HAP1: Vit. C | 0.0001 | | | | | | |

| 5-hmUra | Parental HAP1: Control | Parental HAP1: Vit. C | TET1KO/ TET3KO: Control | TET1KO/ TET3KO: Vit.C | TET2KO/ TET3KO: Control | TET2KO/ TET3KO: Vit. C | TET1KO/ TET3KO: Control |
|---|---|---|---|---|---|---|---|
| TET1KO/TET3KO: Vit. C | 0.0003 | 0.4800 | 0.0009 | 0.6579 | 0.0008 | 0.6486 | 0.0006 |
| TET1KO/TET3KO: Control | 1.0000 | 0.2081 | 1.0000 | 0.2581 | 1.0000 | 0.2664 | |
| TET2KO/TET3KO: Vit. C | 0.1511 | 1.0000 | 0.3414 | 1.0000 | 0.3108 | | |
| TET2KO/TET3KO: Control | 0.9998 | 0.2507 | 1.0000 | 0.3016 | | | |
| TET1KO/TET3KO: Vit.C | 0.1455 | 1.0000 | 0.3316 | | | | |
| TET1KO/TET3KO: Control | 0.9994 | 0.2809 | | | | | |
| Parental HAP1: Vit. C | 0.1048 | | | | | | |

Supplementary Table 3. Tukey's HSD post hoc test results for comparisons of 5-hmCyt, 5-fCyt, 5-caCyt, and 5-hmUra levels in the DNA of HAP1 cells (Parental and with double functional knock-outs of *TET* genes) exposed for 24 hours to 100 µmol/l vitamin C.

| 5-hmCyt fold change | Parental HAP1: Control | Parental HAP1: Vit. C | TET1KO/ TET3KO: Control | TET1KO/ TET3KO: Vit.C | TET2KO/ TET3KO: Control | TET2KO/ TET3KO: Vit. C | TET1KO/ TET3KO: Control |
|---|---|---|---|---|---|---|---|
| TET1KO/TET3KO: Vit. C | 0.7112 | 1.0000 | 0.9589 | 1.0000 | 0.9829 | 1.0000 | 0.9976 |
| TET1KO/TET3KO: Control | 0.8937 | 0.9977 | 0.9994 | 0.9976 | 1.0000 | 0.9976 | |
| TET2KO/TET3KO: Vit. C | 0.7121 | 1.0000 | 0.9592 | 1.0000 | 0.9831 | | |
| TET2KO/TET3KO: Control | 0.9772 | 0.9836 | 1.0000 | 0.9831 | | | |
| TET1KO/TET3KO: Vit.C | 0.7123 | 1.0000 | 0.9592 | | | | |
| TET1KO/TET3KO: Control | 0.9946 | 0.9602 | | | | | |
| Parental HAP1: Vit. C | 0.7153 | | | | | | |

| 5-fCyt fold change | Parental HAP1: Control | Parental HAP1: Vit. C | TET1KO/ TET3KO: Control | TET1KO/ TET3KO: Vit.C | TET2KO/ TET3KO: Control | TET2KO/ TET3KO: Vit. C | TET1KO/ TET3KO: Control |
|---|---|---|---|---|---|---|---|
| TET1KO/TET3KO: Vit. C | 0.0001 | 0.0001 | 0.0001 | 0.0005 | 1.0000 | 0.0441 | 0.4101 |
| TET1KO/TET3KO: Control | 0.0001 | 0.0001 | 0.0001 | 0.0281 | 0.1147 | 0.0001 | |
| TET2KO/TET3KO: Vit. C | 0.0001 | 0.0001 | 0.0001 | 0.0001 | 0.0063 | | |
| TET2KO/TET3KO: Control | 0.0001 | 0.0001 | 0.0001 | 0.0001 | | | |
| TET1KO/TET3KO: Vit.C | 0.0001 | 0.0002 | 0.0639 | | | | |
| TET1KO/TET3KO: Control | 0.0072 | 0.0998 | | | | | |
| Parental HAP1: Vit. C | 1.0000 | | | | | | |

| 5-caCyt fold change | Parental HAP1: Control | Parental HAP1: Vit. C | TET1KO/ TET3KO: Control | TET1KO/ TET3KO: Vit.C | TET2KO/ TET3KO: Control | TET2KO/ TET3KO: Vit. C | TET1KO/ TET3KO: Control |
|---|---|---|---|---|---|---|---|
| TET1KO/TET3KO: Vit. C | 0.0010 | 0.0104 | 0.0001 | 0.2320 | 0.2576 | 0.9301 | 0.0156 |
| TET1KO/TET3KO: Control | 0.9632 | 0.9924 | 0.0318 | 0.9989 | 0.8178 | 0.0002 | |
| TET2KO/TET3KO: Vit. C | 0.0001 | 0.0002 | 0.0001 | 0.0098 | 0.0054 | | |
| TET2KO/TET3KO: Control | 0.1784 | 0.5242 | 0.0003 | 0.9995 | | | |
| TET1KO/TET3KO: Vit.C | 0.8493 | 0.9353 | 0.0502 | | | | |
| TET1KO/TET3KO: Control | 0.3471 | 0.6998 | | | | | |
| Parental HAP1: Vit. C | 1.0000 | | | | | | |

| 5-hmUra fold change | Parental HAP1: Control | Parental HAP1: Vit. C | TET1KO/ TET3KO: Control | TET1KO/ TET3KO: Vit.C | TET2KO/ TET3KO: Control | TET2KO/ TET3KO: Vit. C | TET1KO/ TET3KO: Control |
|---|---|---|---|---|---|---|---|
| TET1KO/TET3KO: Vit. C | 0.0054 | 0.0371 | 0.1713 | 0.0669 | 0.0494 | 0.1473 | 0.1896 |
| TET1KO/TET3KO: Control | 0.6885 | 0.9030 | 1.0000 | 0.9697 | 0.9958 | 0.9983 | |
| TET2KO/TET3KO: Vit. C | 0.9981 | 0.9994 | 0.9990 | 1.0000 | 1.0000 | | |
| TET2KO/TET3KO: Control | 0.9785 | 0.9960 | 0.9976 | 0.9998 | | | |
| TET1KO/TET3KO: Vit.C | 1.0000 | 1.0000 | 0.9768 | | | | |
| TET1KO/TET3KO: Control | 0.7282 | 0.9193 | | | | | |
| Parental HAP1: Vit. C | 1.0000 | | | | | | |

Supplementary Table 4. Tukey's HSD post hoc test results for comparisons of 5-hmCyt, 5-fCyt, 5-caCyt, and 5-hmUra fold change in the DNA of HAP1 cells (Parental and with double functional knock-outs of *TET* genes) exposed for 24 hours to 100 µmol/l vitamin C.

| 5-hmCyt | Control | VIT.C | SCRAMBLED | SCRAMBLED +VIT.C | antiTET2 siRNA |
|---|---|---|---|---|---|
| antiTET2 siRNA +VIT.C | 0.0544 | 0.0001 | 0.0002 | 0.0001 | 0.0001 |
| antiTET2 siRNA | 0.0001 | 0.0001 | 0.0001 | 0.0001 | |
| SCRAMBLED +VIT.C | 0.0001 | 0.0001 | 0.0001 | | |
| SCRAMBLED | 0.3291 | 0.0001 | | | |
| VIT.C | 0.0001 | | | | |
| **5-fCyt** | **Control** | **VIT.C** | **SCRAMBLED** | **SCRAMBLED +VIT.C** | **antiTET2 siRNA** |
| antiTET2 siRNA +VIT.C | 0.9720 | 0.0001 | 0.2114 | 0.0001 | 0.0001 |
| antiTET2 siRNA | 0.0001 | 0.0001 | 0.0001 | 0.0001 | |
| SCRAMBLED +VIT.C | 0.0001 | 0.0001 | 0.0001 | | |
| SCRAMBLED | 0.7795 | 0.0001 | | | |
| VIT.C | 0.0001 | | | | |
| **5-caCyt** | **Control** | **VIT.C** | **SCRAMBLED** | **SCRAMBLED +VIT.C** | **antiTET2 siRNA** |
| antiTET2 siRNA +VIT.C | 0.2810 | 0.0001 | 1.0000 | 0.0001 | 0.0039 |
| antiTET2 siRNA | 0.9677 | 0.0001 | 0.0558 | 0.0001 | |
| SCRAMBLED +VIT.C | 0.0001 | 0.0002 | 0.0001 | | |
| SCRAMBLED | 0.4589 | 0.0002 | | | |
| VIT.C | 0.0001 | | | | |
| **5-hmUra** | **Control** | **VIT.C** | **SCRAMBLED** | **SCRAMBLED +VIT.C** | **antiTET2 siRNA** |
| antiTET2 siRNA +VIT.C | 0.0001 | 0.0462 | 0.0002 | 0.0200 | 0.0001 |
| antiTET2 siRNA | 0.9411 | 0.0001 | 0.9824 | 0.0001 | |
| SCRAMBLED +VIT.C | 0.0001 | 0.9996 | 0.0001 | | |
| SCRAMBLED | 0.7840 | 0.0001 | | | |
| VIT.C | 0.0001 | | | | |

Supplementary Table 5. Tukey's HSD post hoc test results for comparisons of 5-hmCyt, 5-fCyt, 5-caCyt, and 5-hmUra fold change in the DNA of HAP1 TET1KO/TET3KO with reduced TET2 expression.
